# Epigenetic deregulation of IFN and WNT pathways in AT2 cells impairs alveolar regeneration (in COPD)

**DOI:** 10.1101/2023.10.22.563483

**Authors:** Maria Llamazares Prada, Uwe Schwartz, Darius F. Pease, Stephanie T. Pohl, Deborah Ackesson, Renjiao Li, Annika Behrendt, Raluca Tamas, Mandy Richter, Thomas Muley, Joschka Hey, Elisa Espinet, Claus P. Heußel, Arne Warth, Mark Schneider, Hauke Winter, Felix Herth, Charles D Imbusch, Benedikt Brors, Vladimir Benes, David Wyatt, Tomasz P. Jurkowski, Heiko F. Stahl, Christoph Plass, Renata Z. Jurkowska

## Abstract

Chronic lung diseases, including chronic obstructive pulmonary disease (COPD), affect over 500 million people and are a leading cause of death worldwide. A common feature of both chronic and acute lung diseases is altered respiratory barrier integrity and impaired lung regeneration. We hypothesized that alveolar type 2 (AT2) cells, as alveolar epithelial progenitors, will carry molecular alterations that compromise alveolar regeneration in COPD. Sorted AT2 cells from ex-smokers with and without COPD at different disease stages were subjected to RNA sequencing and whole-genome bisulfite sequencing to generate unbiased transcriptome and DNA methylation maps of alveolar progenitors in the lung. Our analysis revealed genome-wide epigenetic changes in AT2 cells during COPD that were associated with global gene expression changes. Integrative data analysis uncovered a strong anti-correlation between gene expression and promoter methylation, suggesting that dysregulation of COPD-associated pathways in AT2 cells may be regulated by DNA methylation. Interferon (IFN) signaling was the top-upregulated pathway associated with the concomitant loss of promoter DNA methylation. Epigenetic regulation of the IFN pathway was validated in both global and targeted DNA demethylation assays in A549 cells. Notably, targeted DNA demethylation of IRF9 triggered upregulation of IFN signaling, mimicking the effects observed in COPD AT2 cells in the profiling data. Our findings suggest that COPD-triggered epigenetic alterations in AT2 cells may impair internal regeneration programs in human lung parenchyma.

## Introduction

Chronic obstructive pulmonary disease (COPD) is the third leading cause of death affecting more than 200 million people worldwide (Safiri et al. 2022). It is a heterogeneous lung disease characterized by irreversible and progressive airflow limitation (Barnes et al. 2015; Rabe and Watz 2017; GOLD 2023). COPD is a risk factor for the development of multiple comorbidities, including lung cancer, depression, hypertension, heart failure, rheumatic disease, muscle wasting and osteoporosis (Stallberg et al. 2016; Yin et al. 2017a). The major risk factor for COPD is cigarette smoke exposure, but the disease is also influenced by ageing, respiratory infections, air pollutants, biomass fuels, as well as genetic and epigenetic factors (MacNee 2009; Barnes et al. 2015; Yin et al. 2017a; Huang et al. 2019; Sakornsakolpat et al. 2019). At the molecular level, COPD pathology is driven by excessive inflammatory responses, increased oxidative stress, degradome alterations and apoptosis of lung progenitor cells (Barnes et al. 2015; Baarsma and Konigshoff 2017; Hikichi et al. 2019). Overall, these altered processes compromise endogenous lung repair and regeneration pathways, leading to structural changes in the small airways, progressive alveolar destruction (emphysema) and ultimately, airflow obstruction (Barnes et al. 2015; Rabe and Watz 2017; Huang et al. 2019). Despite the high economic and health burden caused by COPD, there are no curative therapies, and the clinical approaches are directed solely toward symptom relief and exacerbation control (GOLD 2023). Therefore, understanding the mechanisms regulating human lung regeneration is essential to define novel therapeutic strategies aimed at restoring lung function and respiratory barrier integrity altered in COPD.

Alveolar type-2 (AT2) cells are epithelial progenitors of the lung parenchyma essential for the regeneration of the distal lung and their dysfunction contributes to the development of emphysema in COPD (Liu et al. 2009; Olajuyin et al. 2019; Melo-Narvaez et al. 2020; Ruaro et al. 2021; Lin et al. 2022). AT2 cells produce and secrete surfactant proteins to maintain lung surface tension and play an essential role in maintaining lung homeostasis and repair by proliferating and differentiating into gas-exchanging alveolar type-1 (AT1) cells. AT2 cells in COPD show signs of increased apoptosis (Kosmider et al. 2011), mitochondrial dysfunction (Kosmider et al. 2019a), DNA damage (Kosmider et al. 2019b) and senescence (Tsuji et al. 2006; Tsuji et al. 2010), as well as deregulation of pathways involved in inflammation and lung development, such as interferon (IFN) or WNT/β-catenin signaling (Fujino et al. 2012; Baarsma and Konigshoff 2017; Skronska-Wasek et al. 2017). IFN signaling, which is central to viral immunity, is dysregulated in COPD, in immune and structural cells (Lethbridge et al. 2010; Southworth et al. 2012; Briend et al. 2017; Xu et al. 2019; Booth et al. 2023; Rustam et al. 2023), including AT2 cells (Fujino et al. 2012) and contributes to disease exacerbations. Sustained activation of type I and type III interferon pathways has recently been associated with exacerbated lung pathology in respiratory viral infections due to defective repair caused by disruption of epithelial cell proliferation and differentiation (Major et al. 2020). Similarly, tight control of WNT/β-catenin signaling is essential for AT2 self-renewal and differentiation (Nabhan et al. 2018). Reduced WNT/β-catenin signaling in AT2 cells in COPD is linked to their decreased self-renewal and repair (Baarsma and Konigshoff 2017; Skronska-Wasek et al. 2017; Conlon et al. 2020). Therefore, both signaling pathways are central to alveolar regeneration, and their dysregulation in AT2 cells in a chronically injured environment may drive emphysema progression (Conlon et al. 2020). Thus, understanding the regulatory circuits that drive aberrant gene expression programs in AT2 cells in COPD is of great interest and may identify novel therapeutic strategies to restore endogenous regeneration pathways in COPD.

Epigenetic mechanisms, in particular DNA methylation at CpG sites, are heritable and play a critical role in the regulation of gene expression (Jurkowska and Jurkowski 2019). During differentiation, aging and in response to environmental cues, the epigenome is modified allowing for major changes in transcriptional programs. Alterations in DNA methylation patterns have been implicated in aging, chronic inflammatory diseases, and cancer. In addition, cigarette smoke alters DNA methylation in several clinically relevant samples (Belinsky et al. 2002; Chen et al. 2013; Zeilinger et al. 2013; Wan et al. 2015) and has been associated with altered expression of genes important in COPD pathology (Liu et al. 2010; Vucic et al. 2014; Wan et al. 2015; Yoo et al. 2015; Morrow et al. 2016; Song et al. 2017; Sundar et al. 2017; Clifford et al. 2018; Casas-Recasens et al. 2021; Schwartz et al. 2023). However, previous studies mostly assessed DNA methylation using heterogeneous material with complex cellular composition (e.g., epithelium, blood, or lung tissue) and focused on selected parts of the genome only (mostly gene promoters). To date, no high-resolution, unbiased DNA methylation profiles of purified AT2 cells from COPD lungs are available. Therefore, we set out to profile DNA methylation of AT2 cells at single CpG-resolution across COPD stages to identify epigenetic changes that may drive disease development and progression and combine this with RNA-seq expression profiles. We identified a genome-wide remodeling of the AT2 epigenome in COPD that was associated with global transcriptomic changes. Integrative analysis of the epigenetic and transcriptomic data revealed a strong anticorrelation between gene expression and promoter DNA methylation in COPD AT2 cells, suggesting that aberrant epigenetic changes may drive COPD phenotypes in AT2.

## Results

### Cohort selection criteria and AT2 isolation for an unbiased profiling study in COPD

To identify epigenetic changes associated with disease development and progression, we collected lung tissue from patients with different stages of COPD, which we stratified into three groups based on their lung function data: 1) no COPD (controls) 2) COPD I [stage I, according to the Global Initiative for Chronic Obstructive Lung Disease (GOLD) (GOLD 2023)] and 3) COPD II-IV (GOLD stages II-IV) (**Fig 1A and B**). There were no significant differences between the control group and the COPD patients with respect to gender, age, BMI, and smoking exposure (pack-year), but as expected, the COPD groups could be clearly separated from the control group based on lung function (**Fig 1B**, **Table 1**). We included only ex-smokers to avoid acute smoking-induced inflammation as a confounding factor (van der Vaart et al. 2004). Tissue samples fulfilling the inclusion criteria were cryopreserved and subjected to thorough pathological characterization prior to AT2 isolation and epigenetic profiling (**Fig 1A-C**). This step was critical to avoid potential confounding effects due to the inclusion of patients with additional lung pathologies in the control group as previously documented (Llamazares-Prada et al. 2021). AT2 cells were isolated by fluorescence-activated cell sorting (FACS) from cryopreserved lung parenchyma of three no COPD controls, three COPD I and five COPD II-IV patients as previously described (Fujino et al. 2011; Fujino et al. 2012; Chu et al. 2020) (**Fig 1D-F**). We achieved AT2 purity values ranging between 90-97% as indicated by FACS reanalysis of the sorted cells (**Fig 1F**). The isolated cells were positive for HT2-280, a known AT2 marker (Gonzalez et al. 2010), as confirmed by immunofluorescence (**Fig 1G-H**), validating the identity and high purity of the isolated AT2 populations.

**Figure 1.**
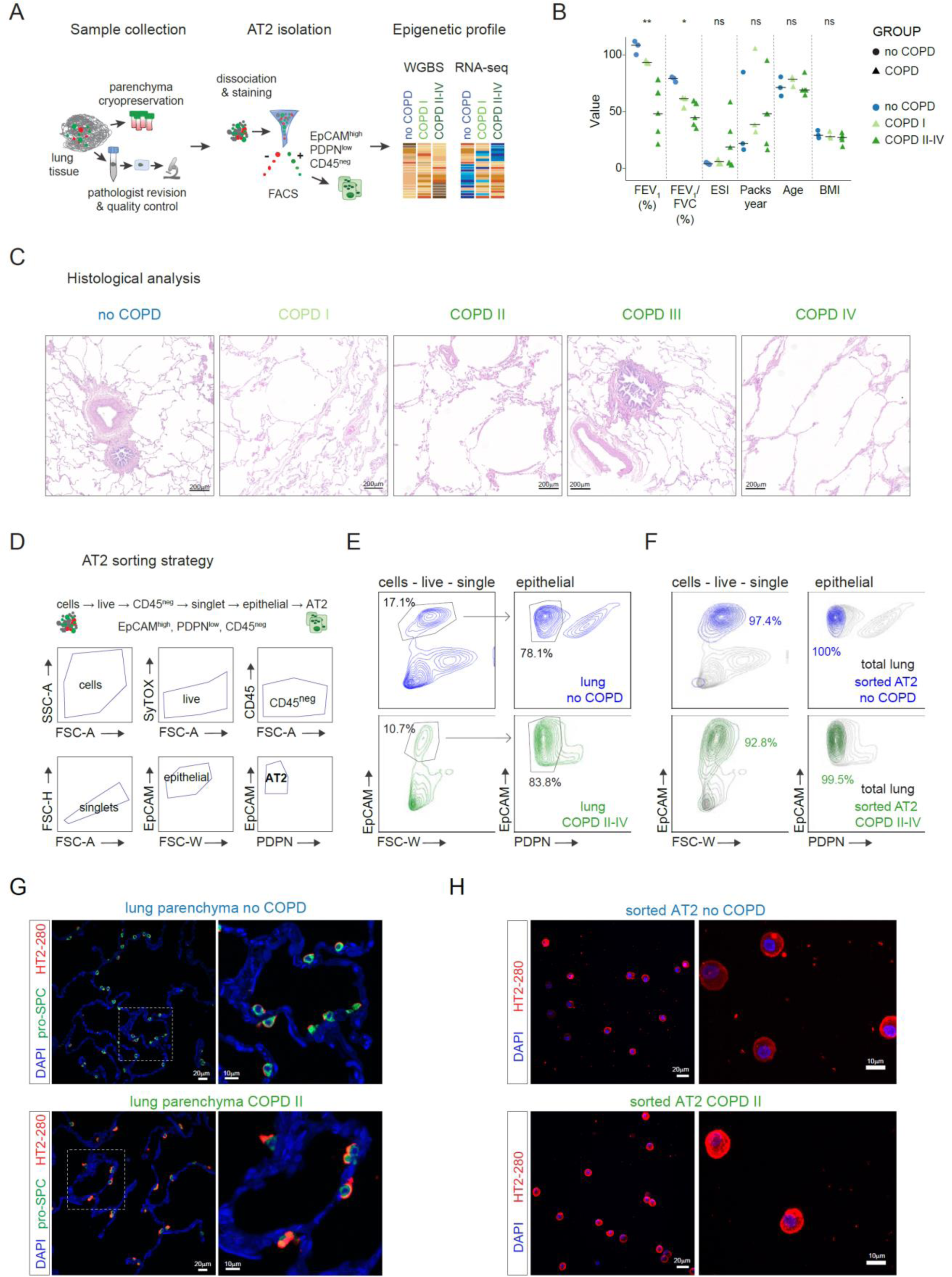
Patient selection and AT2 isolation. **A**. Graphical representation of the experimental approach used in this study, including sample collection, histological characterization, and experimental workflow for profiling. **B**. Patient characteristics. Data points represent values for each donor and horizontal bars the group median. Significant values are indicated **<0.005; *<0.05. One-way ANOVA non-parametric unpaired test was used (Kruskal-Wallis) within the GraphPad Prism software, correcting for multiple comparisons using Dunn’s test. **C**. Representative examples of hematoxylin and eosin (H&E) images of lung tissue samples from samples from the study cohort: no COPD, COPD I, COPD II, COPD III and COPD IV. Scale-bars=200µm. **D**. Sorting strategy employed for isolating AT2 cells from lung tissue. **E**. Representative FACS plots from AT2 sorting for control (blue) and COPD (green) donors. Gates and cell percentages are indicated in the graphs. **F**. Example of FACS plots showing the reanalysis of sorted AT2 cells from control (blue) and COPD (green) donors displayed over total cell lung suspensions from **E** (grey). **G**. Representative immunofluorescence panels of lung parenchyma from control (top) and COPD (bottom) donors showing the presence of AT2 cells (double positive HT2-280, red and pro-SPC, green). Nuclei are counterstained with DAPI, left scale bar=20µm. Right, 3X magnification from marked region on left panels, scale bar=10µm. **H**. Representative immunofluorescence staining images of HT2-280 expression in cytospins from sorted AT2 cells from no COPD (top) and COPD (bottom) donors. Nuclei (blue) were stained with DAPI, left scale bars=20µm; right=10µm.

### The epigenome of AT2 cells is severely altered in COPD

To identify genome-wide DNA methylation changes associated with COPD development and progression in purified AT2 cells, we performed tagmentation-based whole genome bisulfite sequencing (TWGBS) (Wang et al. 2013) (**Fig 1A**). High-quality global DNA methylation profiles at single CpG resolution were generated from 10-20 thousand FACS-sorted AT2 cells (**Table 2**). No global changes in DNA methylation levels were observed between COPD and control samples, when looking at genome wide CpG methylation frequency, but highly methylated CpG sites (> 90%) were significantly underrepresented in COPD II-IV (**Fig 2A**). Unsupervised principal component analysis (PCA) of the most variable CpG sites revealed a separation of COPD II-IV from no COPD on the first principal component (**Fig 2B**), suggesting that variation in DNA methylation across the samples is associated with COPD.

**Figure 2.**
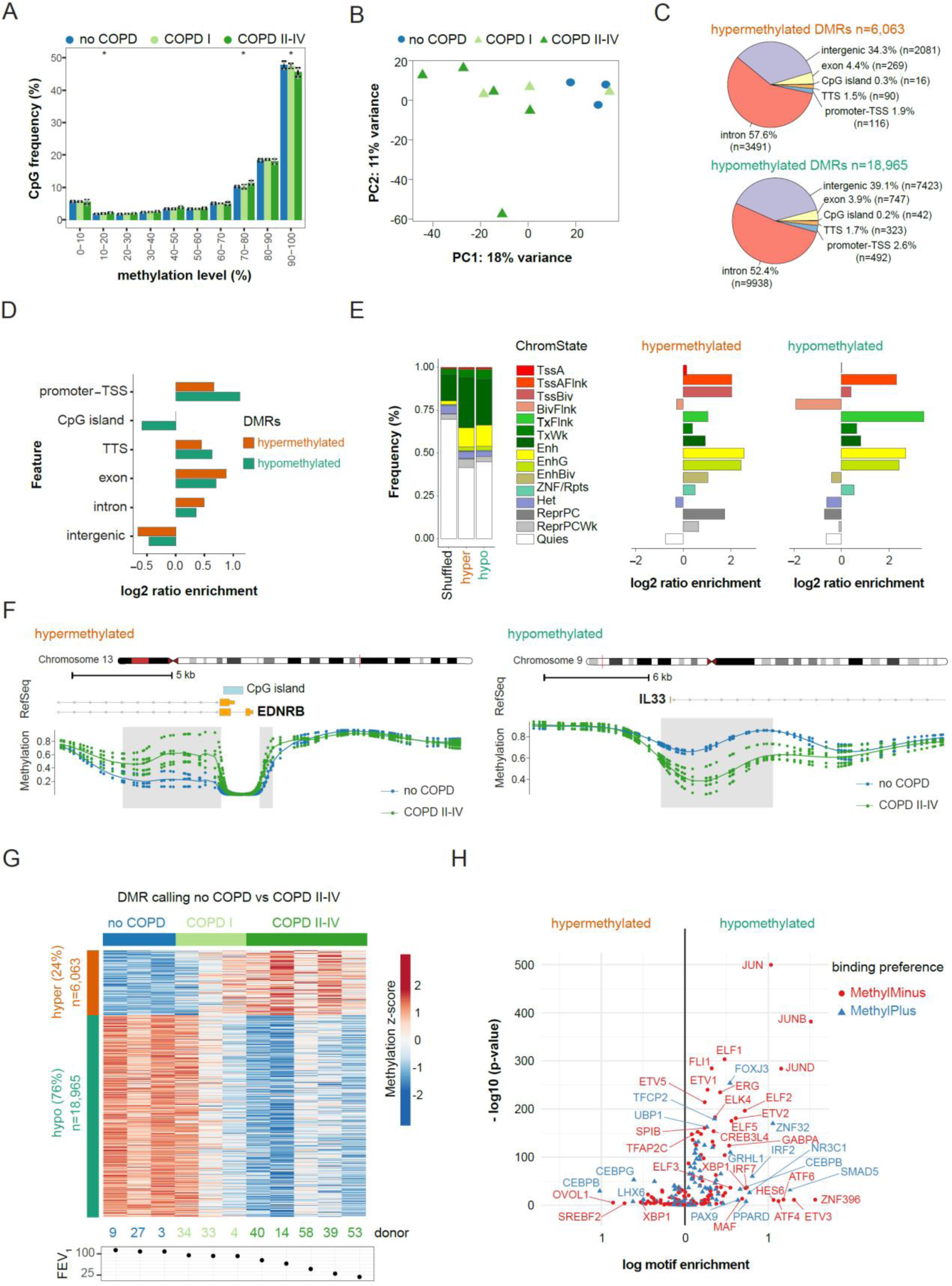
Genome-wide DNA methylation changes occur at regulatory regions in AT2 cells in COPD. Tagmentation-based WGBS data of AT2 cells from no COPD (n=3), COPD I (n=3) and COPD II-IV (n=5) were analyzed at single CpG level **A.** Genome-wide frequency of CpG methylation levels in no COPD, COPD I and COPD II-IV. Methylation levels were binned into deciles. Barplots show the group average and the standard deviations across the samples are indicated. **B**. Principal component analysis (PCA) of methylation levels at CpG sites with > 4-fold coverage in all samples. COPD I and COPD II-IV samples are represented in light and dark green triangles, respectively, and no COPD samples as blue circles. Percentage indicates proportion of variance explained by each component. **C-E**. Genomic feature annotation of differentially methylated regions (DMRs). **C** Distribution of hyper- (top) and hypomethylated (bottom) DMRs at genomic features. TSS, transcription start site TTS, transcription termination site. **D**. Enrichment of genomic features at hyper- (orange) and hypomethylated (green) DMRs compared to the genome wide distribution. **E**. Left panel, distribution of human whole lung tissue specific chromatin states at hypo- and hypermethylated DMRs compared to genome background (shuffled; sampled from regions with matching GC content exhibiting no significant change in methylation). Right panel, chromatin state enrichment relative to the sampled genome background. Abbreviations of chromatin states: Tss= Transcription Start Site; A = Active; Flnk = Flanking; Biv = Bivalent; Enh = Enhancer; Repr = Repressed; PC = Polycomb. **F**. Detailed view of representative hyper- (left, gain) and hypomethylated (right, loss) DMRs (grey box) showing group average (lines) methylation profile and specific CpG methylation levels (dots) of three no COPD (blue) and five severe COPD donors (GOLD II-IV, dark green). RefSeq annotated genes and CpG islands are indicated, if present. **G**. Heatmap of 25,028 DMRs identified in COPD II-IV. Mild COPD (COPD I), which were not used for the DMR calling, are shown within no COPD and COPD II-IV groups. **H**. Enrichment of methylation sensitive binding motifs at hypo- (right) and hypermethylated (left) DMRs. Methylation sensitive motifs were derived from Yin et al (Yin et al. 2017b). Transcription factors, whose binding affinity is impaired upon methylation of their DNA binding motif are shown in red and transcription factors, whose binding affinity upon CpG methylation is increased are shown in blue.

We did not perform differential analysis on individual CpG sites, but for further analysis we focused on differentially methylated regions (DMRs) between no COPD and COPD II-IV. We chose to focus on larger regions rather than individual sites when looking at differential methylation because DNA methylation is spatially correlated and methylation changes in larger regions are more likely to have a biological function. Considering only CpG sites with at least 10% methylation difference and 4x coverage in all samples, we identified 25,028 DMRs between no COPD and COPD II-IV AT2 cells (see Methods for details of DMR calling, **Table 3**). DMRs contained on average 11 CpG sites and were approximately 640 bp in length, indicating that large regions are altered (**Fig S1A-S1B**). Among the identified DMRs, 6,063 regions showed DNA methylation gain in COPD (hypermethylation, 24% of DMRs), while 18,965 displayed DNA methylation loss (hypomethylation, 76% of DMRs), indicating a more permissive chromatin landscape in COPD. Overall, we identified profound, site-specific differences in DNA methylation patterns between primary AT2 cells isolated from COPD patients and non-COPD controls.

### Epigenetic changes in AT2 cells occur in regulatory regions

To better understand the functional role of aberrant methylation in COPD, we investigated the distribution of DMRs across the genome. Both hypo- and hypermethylated DMRs were predominantly located in intronic and intergenic regions (**Fig 2C**). Notably, compared to the genomic background, both types of DMRs were overrepresented at regulatory and gene coding sequences, with hypomethylated DMRs showing the highest enrichment at gene promoters (**Fig 2D**). We further intersected the identified DMRs with known regulatory genomic features in lung tissue annotated by the ENCODE Chromatin States from the Roadmap Epigenomics Consortium (Kundaje et al. 2015). When compared to the genomic background, both hyper- and hypomethylated DMRs were overrepresented at enhancers (Enh) and regions flanking transcription start sites (TSS) of active promoters (TssAFlnk) (**Fig 2E**). In addition, hypermethylated DMRs were overrepresented at Polycomb repressed regions (ReprPC) (**Fig 2E**). The enrichment of DMRs in regulatory regions, including promoters and enhancers, suggests that aberrant DNA methylation in AT2 cells may regulate gene expression in COPD.

To gain insight into cellular processes and pathways affected by aberrant DNA methylation changes in COPD AT2 cells, we linked DMRs to the nearest gene and performed gene ontology (GO) enrichment analysis using the Genomic Regions Enrichment of Annotations Tool (GREAT) (McLean et al. 2010). We identified tube development, epithelial morphogenesis and Wnt signaling among the top categories for hypermethylated DMRs, while genes with hypomethylated DMRs were associated with regulation of reactive oxygen species (ROS) metabolism, protease activity, and negative regulation of MAPK and ERK1/ERK2 cascades (**Fig S1B, Table 4**). Disease-relevant examples of DMRs include two hypermethylated regions upstream and downstream of the TSS of endothelin receptor B gene (EDNRB, 31.3% methylation gain, **Fig 2F**), which could impair the expression of EDNRB in COPD. Furthermore, we found a large hypomethylated DMR region in the first intron of interleukin-33 (IL33, 24,5% methylation loss, **Fig 2F**), an alarmin associated with inflammatory responses and linked to autoantibody production against AT2 cells, exacerbations and disease severity in COPD (Zou et al. 2018; Allinne et al. 2019; Gabryelska et al. 2019). Further DMR examples are provided in **Fig S1C**.

To investigate whether epigenetic dysregulation occurs early in COPD development and to identify methylation changes associated with disease progression, we included TWGBS data from AT2 isolated from COPD I patients (n=3) in the analysis and performed k-means clustering on all identified DMRs using all samples (**Fig 2G**). Consistent with the unsupervised PCA (**Fig 2B**), COPD I samples showed variable methylation changes (**Fig 2G**). Donor 34 displayed a methylation profile similar to the control samples, donor 33 showed an intermediate pattern, while the profile of donor 4 resembled COPD II-IV patients (**Fig 2G**).

### Transcription factor binding sites are enriched at DMRs

Since DMRs were overrepresented at cis-regulatory sites, such as gene promoters and enhancers, we performed motif enrichment analysis to footprint transcription factors (TF) that may mediate the effects of aberrant methylation changes in AT2 cells in COPD (Stadler et al. 2011). Overall, we identified 252 transcription factor binding motifs that were significantly enriched in the differentially methylated regions (**Fig S1D, Table 5**). The specific enrichment in hypomethylated DMRs was obtained for p53/p63, as well as TFs involved in inflammation control, IFN signaling, cell cycle regulation and cell fate commitment, including members of the bZIP (Fra1), homeobox (SIX1) and ETS (ELF3) families (**Fig S1D**). Analysis of hypermethylated DMRs revealed the highest motif enrichment for TFs central for lung development and specification such as NKX-2 and C/EBP (Boggaram 2009; Miglino et al. 2012), suggesting that their binding and function may be affected by DNA hypermethylation in COPD (**Fig S1D**).

To identify TFs reported to change their binding affinity upon methylation of their recognition sites, we used the motifs generated by a systematic methylation sensitivity analysis (Yin et al. 2017b). Interestingly, the motifs of JUN, ELF and ETV proteins, which preferentially bind unmethylated regions, were enriched in hypomethylated DMRs, suggesting that these TFs could bind with higher affinity in COPD and thereby regulate transcription of downstream genes (**Fig 2H**). In contrast, C/EBP TF, which favors binding to methylated CpGs, was enriched in both hypermethylated and hypomethylated DMRs (**Fig 2H**) indicating site specific methylation changes.

Collectively, these results suggest that the aberrant epigenetic makeup of COPD AT2 cells may alter binding of key TFs associated with inflammation, lung development, senescence, apoptosis and differentiation, processes strongly implicated to COPD development and progression.

### Global AT2 transcriptome is severely altered in COPD

The identification of DMRs at cis-regulatory sites and the enrichment of transcription factor motifs in the identified DMRs suggest that changes in DNA methylation may directly impact gene expression in AT2 cells during COPD development. To assess whether epigenetic changes are associated with gene expression changes in AT2 cells in COPD, we performed low-input RNA-seq analysis on FACS-purified AT2 cells matching those used for T-WGBS (**Fig 1A**, **Table 6**). Known AT2-specific genes, including ABCA3, LAMP3 and surfactant genes (SFTPA2, SFTPB and SFTPC) were among the highly expressed genes and were not significantly changed in COPD AT2s (**Fig S2A, Table 7**).

Unsupervised principal component analysis (PCA) on the top 500 variable genes revealed a clear influence of the COPD phenotype in separating no COPD and COPD II-IV samples, as previously observed with the DNA methylation analysis (**Fig 3A**). COPD I samples showed a mixed pattern on PC1 projection and were distributed between no COPD and COPD II-IV with one of the COPD I patients (donor 34) clustering together with no COPD (**Fig 3A**), mirroring DNA methylation data. However, on PC4 projection (**Fig S2B**), COPD I samples separated from the other groups, suggesting a mild-specific expression sub-pattern.

**Figure 3.**
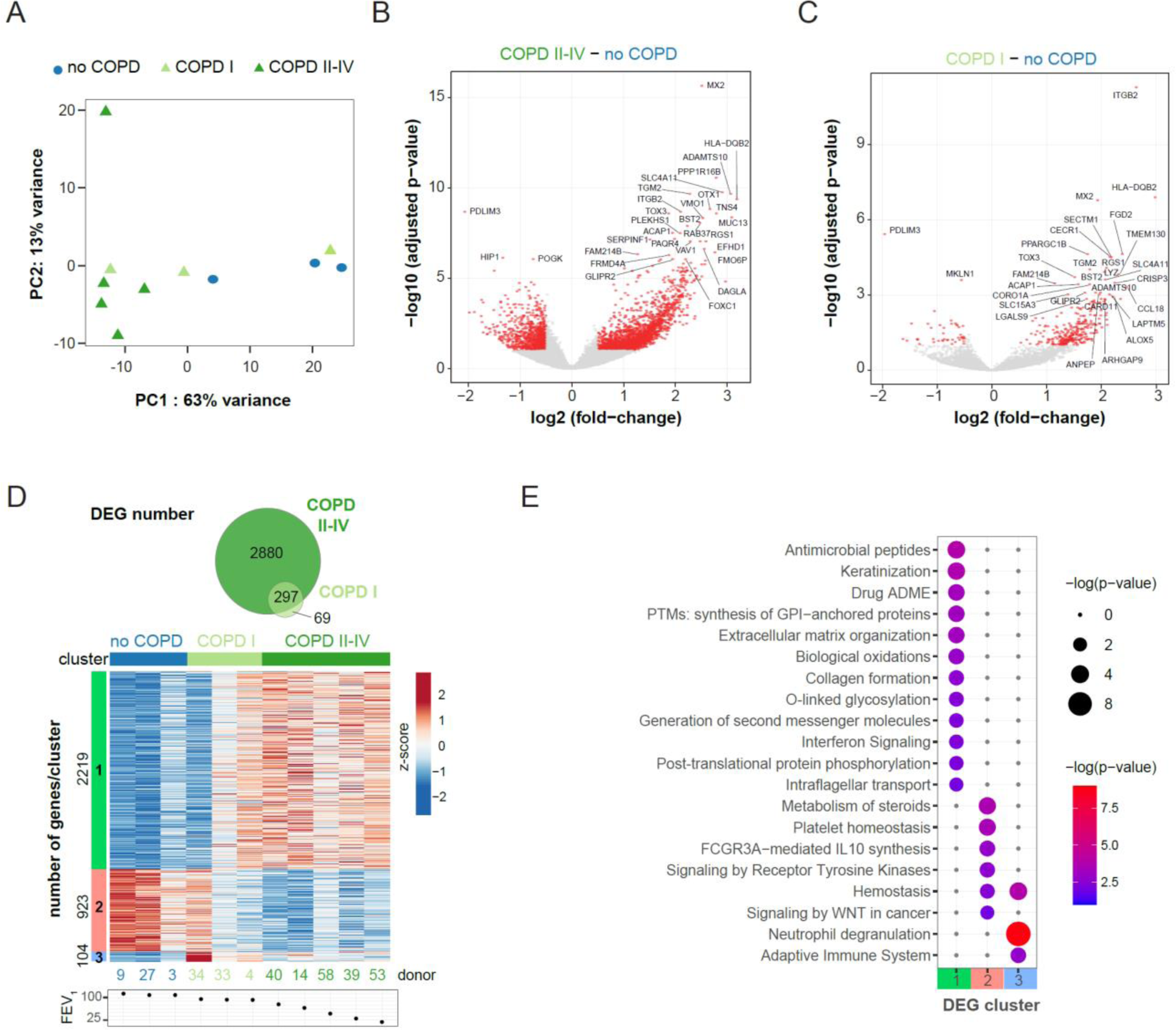
Global transcriptional changes occur in AT2 cells in COPD. **A**. Unsupervised principal component analysis (PCA) of the 500 most variable genes. Percentage indicates proportion of variance explained by each component. **B** and **C**. Volcano plots of differentially expressed genes (DEG) (red dots; FDR of 10% and | log2(fold-change) | > 0.5) in COPD II-IV (**B**) or COPD I (**C**) compared to no COPD. **D**. Top panel, Venn diagram indicating the overlap of DEG in COPD I and COPD II-IV. Bottom, SOM clustering of the DEG displayed in (**B**) and (**C**) using 3 clusters. Donors were sorted according to decreasing FEV_1_ value (indicated below the heatmap). **E.** Metascape enrichment analysis of DEG from each cluster displayed in (**D**). Enrichment p-values are indicated by node size and color. Grey nodes represent non-significant associations.

Differential gene expression analysis identified 2,261 upregulated and 916 downregulated genes in AT2 cells from COPD II-IV group compared to no COPD (**Fig 3B**, |log2 fold-change| > 0.5, at FDR of 10%, **Table 7**), providing the transcriptional signature of COPD. Interestingly, AT2 cells from COPD I already showed transcriptional changes, with 332 upregulated and 44 downregulated genes, providing unique insight into early disease (**Fig 3C**). 297 differentially expressed genes (DEGs) were shared between COPD I and COPD II-IV samples (**Fig 3D**). The most upregulated genes in AT2 cells in COPD were the metalloprotease ADAMTS10 and the Na+-dependent OH− transporter SLC4A11 (**Fig S2D**). ADAMTS10 is involved in microfibril biogenesis (Kutz et al. 2011) and may regulate extracellular growth factor signaling mediated by transforming growth factor beta (TGFβ) and bone morphogenetic proteins (BMP), cytokines essential for proper organ development and tissue architecture (Kutz et al. 2011; Hubmacher and Apte 2015). SLC4A11 upregulation, required for NRF2-mediated antioxidant gene expression (Guha et al. 2017), may be a mechanism to dampen excessive oxidative stress in AT2 cells, a common feature of COPD. PDLIM3 and the potassium channel KCNJ5 were the top two downregulated genes (**Fig S2D**). PDZ-LIM proteins can act as signal modulators, influence actin dynamics and cell migration, regulate cell architecture, and control gene transcription (Krcmery et al. 2010). In the mouse lung, PDLIM5 deficiency has been associated with Smad3 downregulation and emphysema development (Krcmery et al. 2010; Warburton et al. 2013). Our data suggests that PDLIM3 reduction in human AT2 cells from COPD I and COPD II-IV patients may play a role in the development of emphysema.

### COPD relevant pathways are transcriptionally altered in AT2 cells

To further resolve gene expression signatures of COPD states, we performed self-organizing map (SOM) clustering (Wehrens and Kruisselbrink 2018) using combined DEGs from COPD I and COPD II-IV and identified 3 clusters (**Fig 3D**). The largest cluster contained 2,219 genes that were upregulated in both COPD II-IV and in the COPD I donor with the lowest FEV_1_ value (donor 4, FEV_1_=92.1%, **Fig 3D**). Pathway and gene ontology enrichment analysis of this cluster identified processes associated with structural changes in the lung, such as keratinization, extracellular matrix organization, activation of matrix metalloproteinases, and collagen formation (**Fig 3E**, **Table 8**). Antimicrobial peptide signature and interferon signaling were also enriched, suggesting a deregulation of immune and inflammatory signaling in AT2 cells in COPD. Closer examination of the antimicrobial peptide genes revealed that most antimicrobial peptide genes were significantly upregulated in COPD in AT2 cells (**Table 7**). Within the IFN signaling pathway, we observed upregulation of several genes encoding HLA class-I and -II molecules (HLA-F, -DQB2,), GTPases (MX1 and 2), antiviral enzymes (OAS1, 2, 3), immunoproteasome subunits (PSMB8, 9, 10) and transcription factors (IRF5, 7, 9) (**Table 7**). Many of these genes were already upregulated in COPD I samples, suggesting an early activation of the IFN signaling pathway in AT2 cells during COPD pathogenesis. In the second SOM cluster, we identified 923 genes that were downregulated in COPD II-IV and in 2 COPD I donors (**Fig 3D**). DEGs from this cluster are involved in the regulation of cholesterol biosynthesis, tyrosine kinase signaling, WNT signaling, as well as platelet biogenesis and hemostasis (**Fig 3E, Fig S2C**). The smallest cluster contained 104 genes upregulated in COPD with a strong dysregulation already in COPD I donors, and included genes involved in interleukin signaling, neutrophil degranulation and adaptive immune system, emphasizing the dysregulation of inflammatory pathways in AT2 cells as an early event in COPD initiation and progression (**Fig 3E**). To further characterize the DEGs, we performed gene set enrichment analysis (GSEA) and identified overrepresentation of genes related to chronic obstructive pulmonary disease (**Fig S2E**) and epithelial cell differentiation (**Fig S2F**).

Next, we examined pathways involved in AT2 stemness and lung regeneration. We observed a gradual downregulation of axin2, FZD5, LRP2 and LRP6 as well as TCF7L1, known components of WNT signaling (**Table 7**, **Fig S2C**), consistent with dysregulation of WNT signaling in COPD (Baarsma and Konigshoff 2017). In parallel, MCC, a negative regulator of canonical WNT/β-catenin signaling, WNT5B and the cyclin-dependent kinase inhibitors CDKN2A, C and D were upregulated (**Fig S2C**), suggesting a potential loss of repair capacity in AT2 cells in COPD. This was further confirmed when we compared the transcriptional signature of our COPD AT2 with that of alveolar epithelial progenitors (AEPs) identified by Zacharias et al. (Zacharias et al. 2018). We observed a significant negative correlation of gene expression between the DEGs identified in these two datasets (p < 2.2e−16), indicating a decrease in progenitor markers in COPD AT2 cells (**Fig S2G**, **Table 9**).

In addition, we observed a strong upregulation of several keratins (KRT5, KRT14, KRT16, KRT17), mucins (MUC12, MUC13, MUC16, MUC20) and the transcription factor FoxJ1, suggesting a potential dysregulation of AT2 identity and differentiation state in COPD (**Table 7**, **Fig S2F**). Immunofluorescence staining confirmed the presence of KRT5-positive cells in the terminal bronchioles in the distal lung in COPD and identified stretches of cells positive for both KRT5 and HT2-280 signals occurring already in COPD I samples (**Fig S2H**). Collectively, these results indicate a dysregulation of genes involved in the epithelial identity in the terminal bronchioles in COPD, consistent with recent studies (Kadur Lakshminarasimha Murthy et al. 2022; Sauler et al. 2022; Rustam et al. 2023).

### Integrated analysis reveals epigenetically regulated pathways in COPD

DNA methylation is a key mechanism of gene regulation. The similarity of the methylation and gene expression profiles in the PCAs suggested that epigenetic and transcriptomic changes in AT2 cells during COPD might be interrelated (**Fig 2B**, **Fig 3A**). To gain a deeper understanding of the molecular pathways affected by DNA methylation changes in promoters, we assigned identified DMRs to genes in proximity (maximum distance to respective TSS ± 6 kb) (**Fig S3 A-C**). Overall, 755 DEGs, 23.8% of the total, had at least one associated DMR indicating that they might be regulated by promoter proximal methylation (**Fig S3A, Table 10**). We observed a significant overrepresentation of DMRs associated with DEGs compared to non-DEGs (**Fig S3B**, Fisher’s exact test, p-value=2e^-12^). Notably, we observed preferentially a negative relationship between DNA methylation and gene expression, with hypermethylated DMRs mainly associated with downregulated genes and hypomethylated DMRs correlated with upregulated genes (**Fig S3C**). The negative correlation was independent of the location of the DMR relative to the TSS, with hypomethylated DMRs preferentially located downstream of the TSS (**Fig S3C**). Next, we performed Spearman correlation analysis of the promoter DMRs and the corresponding gene expression changes in COPD across all samples. While non-DEGs showed the expected normal distribution, indicating no dependency between promoter methylation and gene expression (**Fig 4A**, blue line), DEGs displayed a bimodal curve enriched at high absolute correlation coefficients (**Fig 4A**, orange line). Among the analyzed DEGs, 76.5% (492) displayed a negative correlation (16.8% of the total DEGs), indicating a possible direct regulation by DNA methylation, while 23.5% (151) showed a positive correlation between gene expression and DNA methylation. We extracted all genes with a Spearman correlation value of > 0.5 or < −0.5 and plotted the methylation differences and expression changes between all COPD and no COPD samples (**Fig 4B**). We observed a clear association between methylation differences and expression changes, with prominent examples including IL33, TMPRSS4, IRF9 and OAS2 (**Table 10**). Enrichment analysis of the negatively correlated genes identified IFN signaling as the top deregulated pathway controlled by promoter proximal methylation (**Fig 4C**, **Table 11**), indicating that aberrant IFN signaling in COPD may be epigenetically regulated.

**Figure 4.**
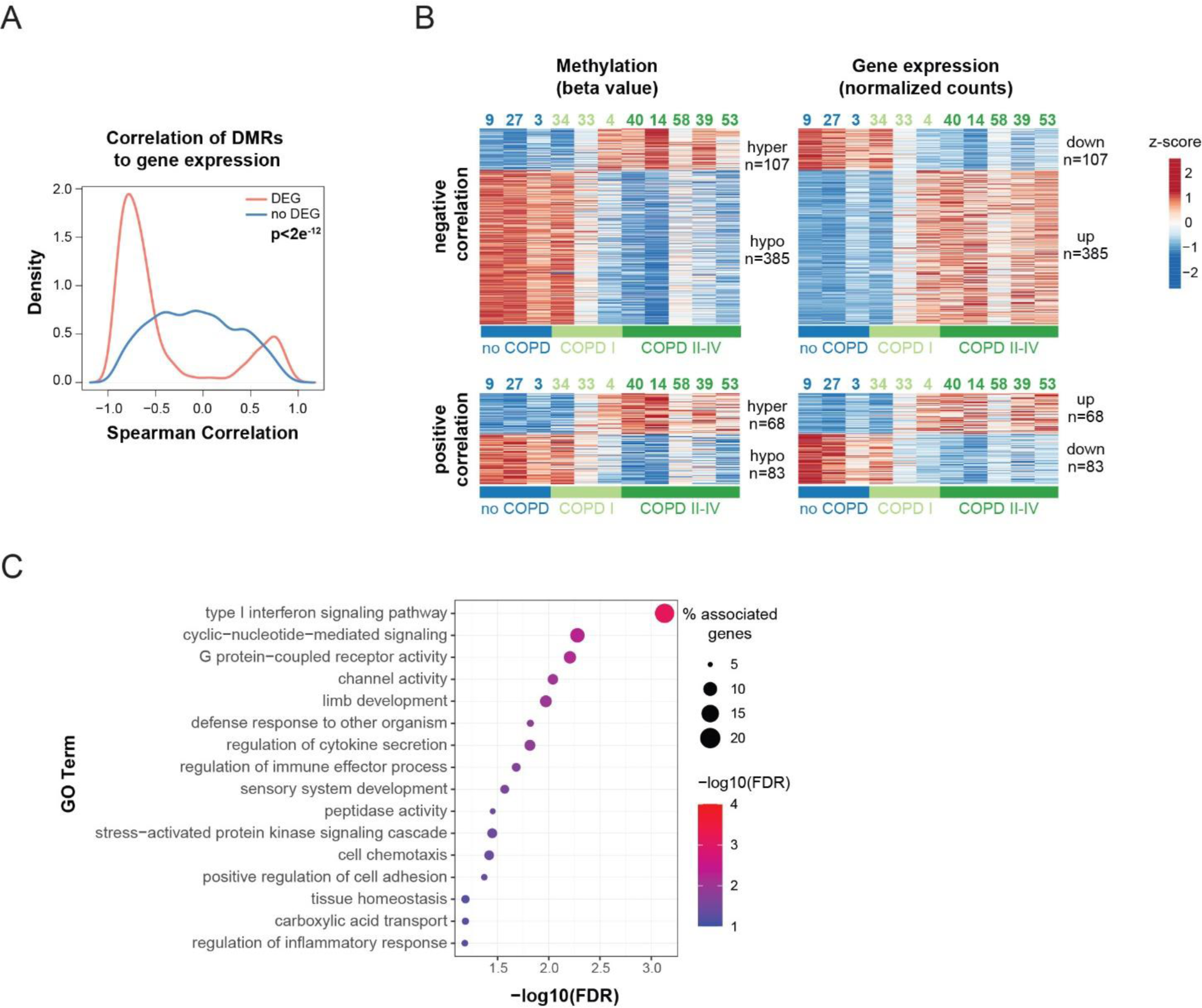
Integrated analysis reveals epigenetic regulation of key AT2 pathways in COPD. **A**. Spearman correlation between gene expression and DMR methylation. DMRs within +/− 6 kb from transcription start site (TSS) were considered. Gene-DMR pairs were split into DEGs (red) and not significantly changed genes (no DEG, blue). **B**. DMR methylation (left, beta-value) and gene expression (right, normalized expression counts) of 492 candidate genes with negative correlation (top, Spearman correlation < −0.5) and 151 genes with positive correlation (bottom, Spearman correlation >0.5) between DNA methylation and gene expression. Values are represented as z-scores. Donor AT2 identifiers are indicated above. **C**. GO-Term over-representation analysis of negatively correlated DEGs. The adjusted p-value is indicated by the color code and the number of associated DEGs is indicated by the node size.

To assess how epigenetic changes in distal regulatory regions (enhancers) relate to transcriptional differences in COPD, we next calculated Spearman correlation between distal DMRs (within 100Kb from TSS) and associated DEGs. We identified several prominent examples with high correlation rates (**Fig S3D**). 147 distal DMRs were associated with 93 genes from WNT/β-catenin pathway (**Fig S3D**), suggesting that DNA methylation may also drive the expression of genes of the WNT/β-catenin family.

### Epigenetic control of IFN and WNT/β-catenin pathways

IFN and WNT signaling are two central pathways associated with AT2 stemness, differentiation and repair (Fujino et al. 2012; Baarsma and Konigshoff 2017; Skronska-Wasek et al. 2017; Nabhan et al. 2018; Conlon et al. 2020; Major et al. 2020). To evaluate whether changes in DNA methylation may regulate the expression of selected key genes of the IFN- and WNT/β-catenin signaling pathways identified from our AT2 study in COPD, we performed a DNA demethylation assay in the alveolar lung cell line A549. A549 cells were treated with increasing doses of the demethylating agent 5-Aza-2’-deoxycytidine (5-AZA). Using mass array, we observed efficient demethylation of the long interspersed nuclear elements (LINEs), indicating successful DNA demethylation upon treatment (**Fig. S4A**). Notably, even cells exposed to low doses of AZA (0.5 µM) showed a clear upregulation of genes from the IFN and WNT/β-catenin signaling pathways (**Fig 5A, S4B**), confirming that their expression may be regulated by DNA methylation.

**Figure 5.**
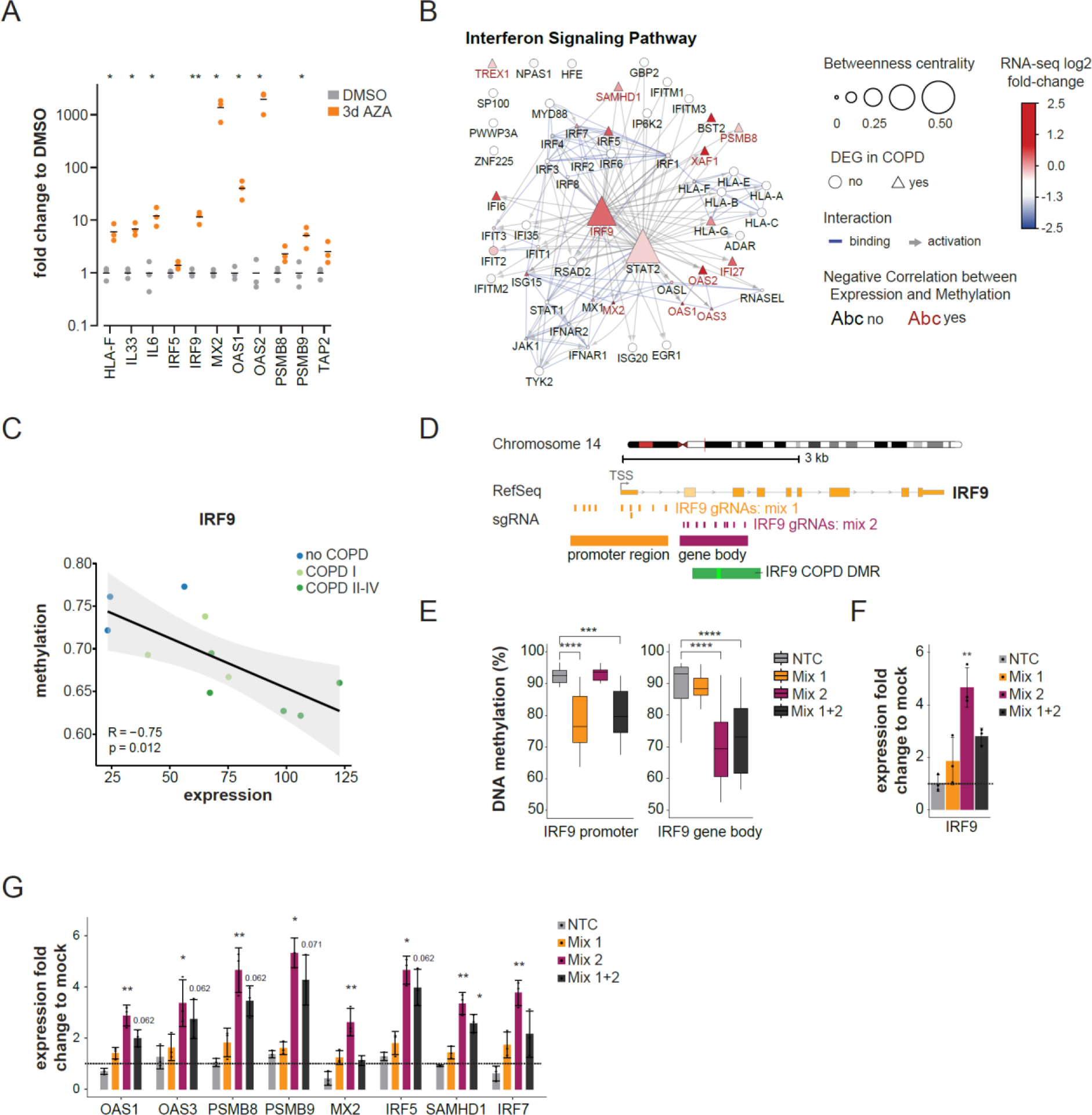
Epigenetic regulation of IFN pathway in AT2 cells. **A**. Relative changes in expression (2^-ΔΔC^_T_) of the indicated IFN pathway genes in A549 cells upon 5-Aza-2’-deoxycytidine (AZA, orange) treatment. Gene expression was measured by RT-qPCR using DMSO treatment as control (grey) and RPLP0 as housekeeping gene. Each point represents the mean of two technical replicates, and bars represent the median of 3 independent experiments (n=3). Paired t-test, FDR corrected using two-stage set-up method of Benjamini, Krieger, and Yekutieli. *p-value < 0.05; **p-value < 0.01. **B.** Cytoscape analysis of components of the IFN pathway identifying IRF9 as a master regulator. Nodes are connected based on their protein-protein interaction annotation in the categories binding (blue line) or activation (grey arrow) in STRING data base. The node size represents the betweenness centrality within the network. DEG are shown as triangles and log2(fold-change) in severe COPD is indicated by the node color. Red labeled nodes exhibit a negative Spearmann correlation (< −0.5) with promoter proximal associated DMRs. **C**. Scatter plots showing correlation between gene expression and methylation of promoter proximal DMRs for IRF9. Each dot represents an individual donor. Dots are color coded according to the disease state. Gene expression is illustrated as normalized counts. Methylation is illustrated as the average beta value of the corresponding DMR. **D**. Detailed view of the IRF9 locus, featuring a core and extended DMR (light and dark green boxes) identified between no COPD and severe COPD. Orange boxes represent the location of individual gRNAs used for targeting the promoter (P, mix 1), and dark purple boxes indicate the location of individual gRNAs targeting the gene body region (GB, mix 2). **E-G** Targeted DNA demethylation of IRF9 using CRISPR-based epigenetic editing in A549 cells. **E.** Box plots displaying percentage DNA methylation at CpG sites across the IRF9 promoter (left) or gene body regions (right) after transfection with the dCas9-VPR-mTet3 demethylating construct and IRF9 targeting gRNA mixes (or non-targeted control; NTC, which contained dCas9-VPR-mTet3 but no gRNA). For each sample, average methylation was calculated per CpG from 3 independent biological replicates and was aggregated into bins containing CpGs from either the promoter or the gene body target regions. Boxes represent the median and IQR of the data, and whiskers represent the full range of non-outlier values. Statistical significance of data was analyzed using the Kruskal-Wallis multiple comparison test, followed by Dunn’s post-hoc analysis comparing each sample against the NTC (total 3 comparisons), and adjusting p-values using the Benjamini-Hochberg correction method (*p-value<0.05; **p-value<0.01; ***p-value<0.001; ****p-value<0.0001). **F-G.** Relative changes in expression (2^-ΔΔC^_T_) of IRF9 (**F**) and a panel of its downstream targets **(G)**. Displayed are the mean fold-changes in gene expression induced by transfection with the dCas9-VPR-mTet3 demethylating construct and IRF9 targeting gRNA mixes (or non-targeted control; NTC) normalized to mock transfection control. Biological replicates data (n=3) is represented by individual data points, and standard deviation by the error bars. Statistical significance of data was analyzed by Kruskal-Wallis multiple comparison tests, followed by Dunns post-hoc analysis test with each sample against the NTC (total 3 comparisons), and adjusting p-values using the Benjamini-Hochberg correction method (p-value<0.05; **p-value<0.01; ***p-value<0.001; ****p-value<0.0001).

However, 5-AZA is a global demethylating agent, and the observed effects may not be direct. Therefore, to further validate the epigenetic regulation of the IFN pathway by DNA methylation, we took advantage of the locus-specific epigenetic editing technology (Jurkowski et al. 2015). Network analysis identified the transcription factor IRF9 as a key master regulator of the upregulated IFN pathway in COPD (**Fig 5B**). Notably, IRF9 upregulation in COPD AT2 cells is associated with a concomitant loss of DNA methylation, suggesting that IRF9 itself may be directly regulated by DNA methylation (**Fig 5C**). This observation was supported by the increased expression of IRF9 (and its downstream target genes) upon 5-AZA treatment of A549 (**Fig 5A**). To investigate the epigenetic regulation of IRF9 expression, we used CRISPR-Cas9-based epigenetic editing to specifically demethylate the IRF9 locus in A549 in a targeted manner. For this, we employed an epigenetic activating domain consisting of the catalytically inactive Cas9 (dCas9) fused to the transcriptional activator VPR and an engineered Tet3 DNA demethylase domain (see Methods for details of the constructs). We targeted the fusion construct to the IRF9 gene using guide RNAs (gRNAs) specific for IRF9 promoter (gRNAs mix 1) or IRF9 gene body (gRNAs mix 2), which contained the demethylated region identified in our profiling data in COPD AT2 (**Fig 5D, Fig. S4E**). Specific demethylation of the targeted regions in the IRF9 locus was validated by amplicon bisulfite sequencing (**Fig 5E, Fig. S4F**). We observed a significant upregulation of IRF9 expression upon specific targeting and DNA demethylation of the IRF9 gene (**Fig 5F**). This effect was not observed in the mock transfected cells (mock), nor in cells transfected with the untargeted dCas9-VPR-TET3 (no gRNA, NCT), where there was also no methylation change (**Fig. 5E-F, Fig. S4F**) confirming the specificity of the assay. Demethylation of the IRF9 region located within the gene body and identified as a DMR in our methylation profiling in COPD AT2 (gRNAs mix 2), showed a stronger demethylation and induction of IRF9 expression than targeting the IRF9 gene promoter itself (gRNA mix 1, **Fig 5E-F**), suggesting the presence of a cis-regulatory element in this region.

Notably, activation of IRF9 by targeted DNA demethylation also resulted in a robust activation of its downstream target genes, including OAS1, OAS3, PSMB8, PSMB9, MX2 and IRF7, demonstrating that demethylation of IRF9 is sufficient to activate IFN signaling pathway (**Fig. 5G**). These experiments confirm the epigenetic regulation of the IFN pathway and validate IRF9 as a master regulator of IFN signaling in alveolar epithelial cells. Taken together, our data suggests an early activation of the IFN signaling pathway in AT2 cells in COPD as a consequence of the epigenetic remodeling of this pathway.

## Discussion

Understanding mechanisms that regulate human lung regeneration is essential to define novel therapeutic strategies aimed at restoring respiratory barrier integrity and lung function impaired in COPD/emphysema and other chronic lung diseases. AT2 cells, as alveolar stem cells, are critical for the maintenance and regeneration of the alveolar epithelium after injury (Nabhan et al. 2018; Zacharias et al. 2018). COPD and emphysema are associated with increased apoptosis, senescence, and decreased regenerative capacity of the alveolar niche (Tsuji et al. 2006; Kosmider et al. 2011; Rustam et al. 2023). However, the molecular regulation of AT2 cell dysfunction in COPD remains poorly understood. In this work, we demonstrate that genome-wide DNA methylation changes occurring in AT2 cells may contribute to COPD pathology by dysregulating key pathways that control inflammation, viral immunity and AT2 regeneration.

Previous studies using various clinical samples provided clear evidence of dysregulated DNA methylation patterns in smokers and COPD patients (Sood et al. 2010; Qiu et al. 2012; Vucic et al. 2014; Wan et al. 2015; Yoo et al. 2015; Busch et al. 2016; Morrow et al. 2016; Sundar and Rahman 2016; Carmona et al. 2018). However, they used low resolution microarray-based approaches, covering a representation of the genome only, and profiled complex samples with heterogenous cellular composition, making direct comparison to our data very difficult. Two studies investigated DNA methylation changes in isolated lung fibroblasts (Clifford et al. 2018; Schwartz et al. 2023), but the epigenetic dysregulation of AT2 cells across COPD stages remained uncharted.

Using high-resolution epigenetic profiling, we uncovered widespread alterations of DNA methylation landscape in AT2 cells in COPD that were associated with global gene expression changes. Consistent with our earlier methylation data from COPD lung fibroblasts (Schwartz et al. 2023), epigenetic changes in AT2 were enriched at cis-regulatory regions, including enhancers and promoters, indicating that they may drive aberrant transcriptional programs in COPD. This was further supported by the strong anticorrelation between DNA methylation and gene expression, with more than 500 dysregulated genes showing corresponding DNA methylation changes in promoters. WNT and IFN signaling, two central pathways orchestrating the regenerative capacity of AT2 cells (Nabhan et al. 2018; Zacharias et al. 2018; Katsura et al. 2020) were enriched for genes with anticorrelated DNA methylation signatures in promoters and distal regions, respectively. Hence, epigenetic dysregulation of key pathways involved in AT2 proliferation, renewal and differentiation may be one of the mechanisms contributing to the lack of renewal of alveolar cells in COPD and emphysema.

Chronic inflammation has long been recognized as a pathogenic driver that exacerbates COPD phenotypes, including airway remodeling and progressive alveolar destruction (Barnes 2019; Mehta et al. 2020; Booth et al. 2023). Notably, the inflammatory processes in COPD persist long after smoking cessation (Shapiro 2001), suggesting an epigenetic regulation, yet the molecular mechanisms driving inflammation and lung tissue destruction in COPD are not fully understood. We observed upregulation of several IFN genes in AT2 in COPD, consistent with a previous study (Fujino et al. 2012). Upregulation of the IFN genes was correlated with a concomitant loss of DNA methylation at their promoters, indicating their epigenetic control in COPD. Notably, epigenetic remodeling of IFN-signaling genes occurred already in early stages of COPD (in COPD I), suggesting that it may contribute not only to disease exacerbation but also to its development. Interferons are essential for antiviral host defense. They are induced upon viral recognition through the binding of interferon-responsive transcription factors (IRFs) and coordinate the expression of IFN-stimulated genes (ISGs) in the infected and neighboring cells (Barrat et al. 2019). Dysregulated IFN signaling has been reported in smokers and COPD patients and has been associated with impaired antiviral immunity, increased susceptibility to infections and disease exacerbations (Hilzendeger et al. 2016; Hsu et al. 2016; Wu et al. 2016; Garcia-Valero et al. 2019; Mehta et al. 2020).

In addition to its pro-inflammatory and antiviral roles, IFN also has anti-proliferative and apoptotic effects (Parker et al. 2016). In particular, sustained activation of IFN induces lung tissue remodeling and destruction. Targeted pulmonary overexpression of IFN-γ in mice causes inflammation and leads to emphysema development via induction of proteolytic enzymes (e.g., MMP-12) (Wang et al. 2000). IFN-γ is also a potent inducer of DNA damage and apoptosis in airway and AT2 cells (Zheng et al. 2005). Recently, a direct effect of IFN on alveolar epithelial repair was demonstrated, as prolonged IFN production reduced AT2 proliferation and differentiation during recovery from influenza infection (Major et al. 2020). Similarly, IFN has also been implicated in impaired distal airway regeneration in COPD, as progenitor generation of terminal airway-enriched secretory cells (TASCs) was suppressed by IFN-γ signaling, which was increased in the distal airways of COPD patients (Rustam et al. 2023). Our data showing epigenetically driven upregulation of IFN-signaling in AT2 cells may provide a mechanistic explanation for the reduced repair capacity of alveolar epithelial cells in COPD. Indeed, we also observed a strong downregulation of alveolar epithelial progenitor (AEP) markers (Zacharias et al. 2018) in AT2 cells from COPD patients, indicative of their impaired regenerative potential.

Currently, it is unclear how cigarette smoking leads to changes in DNA methylation patterns in AT2 and how the epigenetic changes translate into biological phenotypes in COPD. We have identified IRF9 as a key epigenetically regulated master regulator of IFN activation. IFNs can induce extensive remodeling of the epigenome, including histone marks, through binding of the IRFs to gene regulatory elements and regulation of chromatin accessibility at promoters and enhancers (Park et al. 2017; Kamada et al. 2018). They recruit chromatin-remodeling enzymes leading to the transcriptional activation of ISGs (Barrat et al. 2019). IFN-induced epigenetic changes can persist beyond the period of IFN stimulation, conferring transcriptional memory and sustained expression of ISGs. This “epigenetic priming” may mediate the sensitivity of AT2 cells to subsequent environmental exposures, contributing to impaired antiviral responses and reduced alveolar regeneration in COPD. Treatment with a demethylating drug and targeted epigenetic editing demonstrated the ability to modulate the expression of IFN-stimulated genes through demethylation of IRF9. Therefore, effective modulation of IFN signaling to restore the robust induction of ISGs upon injury/exposure, followed by timely downregulation of IFN responses to allow efficient alveolar epithelial repair, may provide protection against disease exacerbations and enhance the regenerative capacity of AT2 cells in COPD patients.

DNA methylation in regulatory regions can modulate the binding of transcription factors to DNA (Stadler et al. 2011), thus methylation profiling allows the identification of transcriptional regulators potentially mediating the epigenetic changes. In line with this, in the identified DMRs we detected a significant enrichment of binding sites of transcription factors associated with lung development, apoptosis, senescence, inflammation and differentiation, processes critical for tissue repair and regeneration that are altered in COPD. Binding sites for the lineage factor NKX2 and TEAD TFs were enriched in hypermethylated DMRs, suggesting that their changed binding could impact cell fate determination in AT2 cells during lung regeneration (Little et al. 2021). Moreover, TP53 binding sites were overrepresented in hypomethylated DMRs in COPD, which could potentially explain the increased apoptosis of AT2 cells reported in COPD lungs (Kosmider et al. 2011). Consistent with this hypothesis, recent studies demonstrated that p53 mediates apoptosis of cycling cells mobilized during tissue repair in lungs exposed to prolonged inflammation during COVID-19 and influenza infection (Katsura et al. 2020; Major et al. 2020). Taken together, our data suggests that NKX2 and TP53, whose binding sites are enriched in DMRs, may mediate some of the downstream biological effects in COPD AT2 and contribute to COPD phenotypes. Future work is needed to delineate and experimentally validate the target genes directly bound and regulated by these transcription factors in AT2.

Not much is known about the correlation of DNA methylation with disease severity. 13 genes with altered methylation patterns have been identified in the lung tissue of COPD GOLD I and II patients compared to non-smoking controls (Casas-Recasens et al. 2021). Our previous study revealed that genome-wide DNA methylation changes are present in lung fibroblasts from COPD I patients compared to controls with matched smoking history, demonstrating that epigenetic changes occur early in COPD development and may provide a sensitive biomarker for early disease detection. Interestingly, we observed heterogeneous methylation profiles within the mild COPD group in AT2 cells, despite almost identical lung function data of the three COPD I donors, suggesting that epigenetic profiling may provide an additional layer of information for differentiating mild COPD patients and disease progression. However, our cohort included only 3 COPD I donors and we did not have any follow-up data on the patients, so future large-scale profiling of early disease in an extended patient cohort will be crucial for a better understanding of early disease and its progression trajectories.

Overall, our findings suggest that rewiring the epigenomic landscape in COPD in AT2 cells to revert aberrant transcriptional programs may have the potential of restoring the internal regenerative programs lost in the disease.

## Methods

### Patient samples

Human lung tissue samples were obtained from the Thoraxklinik Heidelberg (Germany) from patients undergoing lung surgery due to primary squamous cell carcinomas (no COPD and COPD, GOLD I-II) or lung resection (COPD, GOLD III-IV). All patients gave written informed consent and remained anonymous in the context of this study. The protocol for tissue collection was approved by the ethics committees of the University of Heidelberg (S-270/2001) and followed the guidelines of the Declaration of Helsinki.

Strict patient inclusion criteria were established at the beginning of the tissue collection to ensure the best possible matching of control and disease groups in terms of age, BMI, gender, smoking status (ex-smokers) and smoking history. To avoid acute inflammation due to smoking all the collected donors have ceased smoking for at least 3-months. None of the donors received chemotherapy or radiation within 4 years before surgery. In addition, lung function results, based on the forced expiratory volume in 1 second (FEV_1_) as well as the FEV_1_/forced vital capacity ratio (FEV_1_/FVC ratio), quantitative emphysema score index based on chest CT as well as medical history were collected for each patient for the best possible characterization of the included samples. All tissue samples underwent strict evaluation by an experienced lung pathologist, who confirmed the absence of tumors in all the samples as well as lack of extensive fibrosis and emphysema in the control group (**Fig 1C**). Besides, COPD relevant phenotypes, such as emphysema, airway thickening, and immune infiltration were evaluated. In total, 3 smoker controls, 3 mild COPD donors (GOLD I) and 5 moderate-to-severe COPD donors (GOLD II-IV) were profiled (**Fig 1A-C**, **Table 1**). Importantly, 3 of the profiled samples from donors with severe COPD (HLD39, 53 and HLD58) came from lung resection, thus representing tissue without cancer background, an important control, ensuring that the observed changes are also present in cancer-free material.

### Emphysema index determination

Lung and emphysema segmentation were performed calculating the emphysema score index (ESI) from clinically indicated preoperative CT scans taken with mixed technical parameters. After automated lung segmentation using the YACTA-software, a threshold of −950 HU was used with a noise-correction range between −910 and-950 HU to calculate the relative amount of emphysema in % of the respective lung portion (Lim et al. 2016). While usually global ESI was measured, only the contralateral non-affected lung side was used if one lung was severely affected by the tumor.

### FFPE and H&E

Representative slices from different areas of the tissue samples were taken and fixed O/N with 10% neutral buffered formalin. Next, fixed slices were washed and kept in 70% ethanol. Sample dehydration, paraffin embedding, and H&E staining was performed at Morphisto (Morphisto GmbH, Frankfurt, Germany). Two 4 µm thick sections were cut per sample on a Leica RM2255 microtome with integrated cooling station and water basin and transferred to adhesive glass slides (Superfrost Plus, Thermo Fisher Scientific). Subsequently, sections were dried to remove excess water and enhance adhesion. Anonymized H&E-stained slides were evaluated by an experienced lung pathologist at the Thorax Clinic in Heidelberg.

### Cryopreservation of lung parenchyma, airway, and vessel enriched fractions

Lung specimens were transported in CO_2_ independent medium (ThermoFisher Scientific) supplemented with 1% BSA (Carl Roth), 1% penicillin/streptomycin (Fisher Scientific) and 1% amphotericin B (Sigma) and processed as previously published (Llamazares-Prada et al. 2021; Pohl et al. 2023). Briefly, tissue pieces were inflated with cold HBSS (Fisher Scientific) supplemented with 1% BSA (Carl Roth), 2mM EDTA (Fisher Scientific), 1% Amphoterin B and 1% penicillin/streptomycin (Fisher Scientific) (later referred as HBSS^++++^) and exemplary samples of the lung piece were collected for histological analysis (see above). Pleura, airways and vessels were removed, and the airway and vessel-free parenchyma was further minced 4 x 4 mm pieces and cryopreserved in high glucose DMEM GlutaMAX^TM^ (Thermo Fisher Scientific) containing 10% DMSO (Carl Roth) and 20% FBS (GE Healthcare). For long-term storage, tubes were kept in liquid nitrogen.

### Tissue dissociation, viability check and FACS sorting

Cryo-tubes containing lung parenchyma were thawed, washed, and minced into smaller pieces prior to mechanical and enzymatic dissociation as previously reported (Llamazares-Prada et al. 2021). Tissue pieces were dissociated into single cell suspensions with the human tumor dissociation kit following manufacturer instructions (Miltenyi Biotec). Digestion was stopped with 20% fetal bovine serum (FBS, Gibco, 10270106) and single cells were washed and collected by sequential filtering through 100 µm, 70 µm and 40 µm cell strainers (BD Falcon). Cells were spun down, incubated for 5 minutes in ACK lysis buffer (Sigma Aldrich) at room temperature for red blood cell lysis. After two washes with HBSS^++++^ to neutralize cell lysis buffer, Fc receptors were blocked with human TruStain FcX (Biolegend) for 30 minutes on ice. Immune and epithelial cells were labelled using EpCAM, PDPN and CD45 antibodies (Anti-human CD326 (EpCAM-PE, 12-9326-42, Affymetrix eBioscience), Anti-human podoplanin (PDPN)-Alexa Fluor 647 (337007, BioLegend), Anti-human CD45-PerCP-Cy5.5 (45-9459-42, Affymetrix eBioscience), CD45-APC Cy7 (BioLegend, 304014), or CD45-Bv605 (564047, BD)) for 30 minutes in the dark at 4°C following manufacturer instructions. Stained samples were washed, resuspended in HBSS^++++^ and added to FACS tubes with 40 µm cell strainer caps. To discriminate between live and dead cells, we used SyTOX blue as recommended by manufacturer (Thermo Fisher Scientific). We sorted live, single-cell gated, EpCAM^high^ PDPN^low^ cells as previously published (Fujino et al. 2011; Fujino et al. 2012; Chu et al. 2020) using a FACS Aria IIu cell sorter. Sorted epithelial cells were manually counted, aliquoted, spun down and flash-frozen in liquid nitrogen. Cell pellets were kept at −80C until the full cohort was collected for subsequent RNA-seq and TWGBS to avoid batch-effects. The remaining cells were fixed for immunofluorescence studies. FlowJo software (Tree Star) was used to analyze the FACS results.

### Immunofluorescence

FFPE lung tissue samples were cut in 4 µm thick sections and added to Superfrost Plus slides (Thermo Fisher Scientific), deparaffinized and rehydrated by immersing in Xylene and gradual decreasing solutions of ethanol as published (Llamazares-Prada et al. 2021). Antigen retrieval was conducted at 99°C for 20 minutes in a pressure cooker containing citrate buffer (10 mM citric acid, 0.05% Tween 20, pH 6.0) and allowed to cool down for 90 minutes at room temperature. Tissue samples were surrounded with a hydrophobic pen (abcam) and permeabilized with 1% Triton-X100 (Carl Roth) in PBS 1X for 20 min at RT without shaking. Slides were washed with PBS 1X and endogenous peroxidase was quenched with 3% H_2_O_2_ for 1h at RT following manufacturer’s instructions (Tyramide SuperBoost kit (TSB), Thermo Fisher Scientific). Slides were washed with PBS 1X, blocked with 10% goat serum (TSB kit, Thermo Fisher Scientific) for 1h at RT prior to overnight incubation with HT2-280 (IgM mouse monoclonal, 1:30 dilution, Terrace Biotech), pro-SFTPC (IgG rabbit polyclonal, 1:200 dilution, abcam ab196677) and/or KRT5 (IgG rabbit monoclonal, 1:200 dilution, abcam ab52635) primary antibodies at 4°C in incubation buffer (TSB kit, Thermo Fisher Scientific). Mouse IgM and rabbit IgG were used as controls. Tissues were washed thoroughly with PBS 1X before incubating 1h at RT with secondary antibody solution from TSB kit containing secondary goat-anti-rabbit IgG coupled to HRP in 10% normal goat serum. Secondary goat-anti-mouse-IgM-AF568 (1:500 dilution, Thermo Fisher Scientific) was added to the mix. Slides were washed with PBS 1X before signal amplification was performed following manufacturer’s instructions. Briefly, slides were incubated for 3 minutes with tyramide working solution (inactive tyramide connected to AlexaFluor 488) and the reaction was blocked with working stop solution for 1 minute at RT. Finally, tissue slides were mounted with ProLong Antifade reagent containing DAPI (Thermo Fisher Scientific). Microscope slides were left to dry overnight before imaging. Imaging of cells was conducted at the ZMBH imaging facility (Heidelberg) using the Zeiss LSM780 confocal fluorescent microscope.

### DNA extraction and TWGBS

Genomic DNA was extracted from 1-2×10^4^ sorted alveolar epithelial cells isolated from cryopreserved lung parenchyma from 11 different donors using QIAamp Micro Kit (Qiagen, Hilden, Germany) following manufacturer’s protocol, with an additional RNase A treatment step (Qiagen, Hilden, Germany). TWGBS was essentially performed as described previously (Wang et al. 2013; Schwartz et al. 2023) using 25-30 ng genomic DNA as input. Four sequencing libraries were generated per sample using 11 amplification cycles. Equimolar amounts of all four libraries were pooled and sequenced on two lanes of a HiSeq2500 (Illumina, San Diego, California, US) machine using NGX Bio service (San Francisco), with 100 bp, paired end reads.

### TWGBS read alignment and methylation quantification

Read alignment and methylation quantification were performed as described (Schwartz et al. 2023). Briefly, the MethylCtools pipeline was modified for TWGBS data and used for whole genome bisulfite sequencing mapping (Hovestadt et al. 2014). Adaptor sequences were trimmed using Trimmomatic (Bolger et al. 2014). For alignment, cytosines in the reference and read sequences are converted to thymines prior to alignment. The reads were aligned to the transformed strands of the hg19 reference genome using BWA MEM, then reverted to their original states. Duplicate reads were marked using Picard MarkDuplicates (http://picard.sourceforge.net/). For methylation calling, the cytosine frequency was used to determine methylated CpGs, whereas cytosine to thymine conversion indicated unmethylated CpGs. Only bases with a Phred-scaled quality score of ≥ 20 were considered, excluding the 10 bp at the ends of the reads and CpGs on sex chromosomes.

### DMR calling

DMR calling between severe COPD and no COPD was performed as described (Schwartz et al. 2023). Shortly, The R**/**Bioconductor package bsseq was used to detect differentially methylated regions (Hansen et al. 2012). The data were first smoothed using the Bsmooth function, and only CpG sites with at least 4x coverage were used for further analysis. A t-statistic was calculated between the two groups using the Bsmooth.tstat function. Differentially methylated regions (DMRs) were identified by selecting the regions with the 5% most extreme t-statistics, filtering for regions with at least 10% methylation difference and containing at least 3 CpGs. A non-parametric Wilcoxon test was applied to remove potentially false positive regions, since the t-statistic is not well-suited for not normally distributed values, as expected at very low/high (close to 0% / 100%) methylation levels. A significance level of 0.1 was used without further FDR correction. Additionally, DMRs which were < 5 kb apart from each other were stitched together if they had the same direction of methylation change (hyper/hypo) and the resulting average methylation level within the DMR did not drop below 10%.

### DMR downstream analysis

The R/Bioconductor package methylkit was used to generate the CpG methylation frequency distribution and PCA plot excluding CpGs with less than 4 fragment coverage and the CpGs with 0.1% highest coverage (Akalin et al. 2012). Genome feature annotation and known transcription factor motif enrichment analysis was performed using the HOMER functions annotatePeaks.pl and findMotifsGenome.pl (Heinz et al. 2010). Genome tracks were plotted using the R/Bioconductor package Gviz (Hahne and Ivanek 2016). Roadmap chromatin states were obtained for lung tissue (E096) (Roadmap Epigenomics et al. 2015). DMRs within 100 kb distance were assigned to the next gene and subjected to gene ontology enrichment analysis using GREAT (McLean et al. 2010). To define significant associations with pathways, we used the default settings of the GREAT tool, which are as follows: FDR < 0.05 in both binominal and hypergeometric tests and minimum region-based fold enrichment of 2.

### RNA isolation and RNA-seq

RNA was isolated from flash frozen pellets of 2×10^4^ sorted AT2 cells using the Arcturus Picopure RNA Isolation kit (Thermo Fisher Scientific, KIT0204) following manufacturer’s instructions. DNA was removed by on column Dnase treatment (Qiagen) before elution with nuclease-free water (Thermo Fisher Scientific). RNA concentration and integrity were measured using the RNA Pico 6000 Assay Kit of the Bioanalyzer 2100 system (Agilent Technologies, Santa Clara, CA), and only samples with RIN > 8 were included in RNA-seq. Low-input, stranded mRNA (poly-A enriched) libraries were manually prepared at the Genomics core facility (GeneCore) at EMBL, Heidelberg (Germany). Obtained libraries were pooled in equimolar amounts. 1.8 pM solution of each library was pooled and loaded on the Illumina sequencer NextSeq 500 High output and sequenced uni-directionally, generating ∼450 reads per run, each 75 bases long.

### Alignment and transcript abundance quantification

The raw sequence data was processed using Trimmomatic v0.36, a flexible read trimming tool for Illumina sequence data (Bolger et al. 2014). The parameters for Trimmomatic were set to remove adapters (**ILLUMINACLIP**), trim low-quality bases from the start (**LEADING:3**), the end (**TRAILING:3**), and perform a sliding window trimming (**SLIDINGWINDOW:4:15**), with a minimum length of the reads set to 36 bases (**MINLEN:36**). The trimmed reads were then aligned to the reference genome hg19 and transcriptome (Ensemble release 87) using the Spliced Transcripts Alignment to a Reference (STAR) aligner (Dobin et al. 2013). The parameters were set to filter out alignments with high mismatch rates and multi-mapping reads. Quality reports were generated using Qualimap’s RNA-seq QC module (Okonechnikov et al. 2016). Duplicate reads were marked using the **bam dedup** command from bamUtil package (https://github.com/statgen/bamUtil). The counting was performed using the **featureCounts** function from the Subread package (Liao et al. 2014). The parameters were set to count reads in reverse stranded libraries (**-s 2**), ignore duplicate reads (**--ignoreDup**)

### Differential gene expression analysis

The DESeq2 package (Love et al. 2014) was used to read the count table and to create a DESeqDataSet object from the count data and meta data. The count data was subset to include only autosomes and only lincRNA and protein coding genes. Exploratory analysis was performed to visualize the distribution of raw or rlog transformed counts, and to run PCA. The DESeq2 package was also used to perform differential gene expression analysis. The results were filtered to include only significant hits (adjusted p-value < 0.1 and absolute log2 fold change > 0.5). Gene set over-representation analysis was carried out using the Metascape online tool (Zhou et al. 2019). For this analysis, the background was defined as all expressed genes. Significantly differentially expressed genes were categorized based on self-organizing map clustering. The following settings were applied: p-value cutoff of 0.01, and minimum enrichment of 1.5. Enrichment was performed using the Reactome Gene Set. This analysis helped to identify the key functional categories that were overrepresented in the respective cluster. Selected over-represented terms from Metascape analysis were visualized using the Bioconductor package clusterProfiler (Wu et al. 2021).

### Integrated analysis

Cytoscape was used to analyze negatively correlated DMR DEG pairs. ClueGO (v2.5.6) analysis was conducted using all DEG associated with a promoter proximal DMR and correlation coefficient < −0.5 (Bindea et al. 2009). The following settings were used: statistical test used = Enrichment (Right-sided hypergeometric test), correction method used = Benjamini-Hochberg, Min GO Level = 4, Max GO Level = 10, Kappa Score Threshold = 0.4. Next genes associated with the top enriched term “type I interferon signaling pathway” were extracted und a gene interaction network was built using the Cytoscape plugin CluePedia (Bindea et al. 2013).

### A549 cell culture and 5-Aza-2’-deoxycytidine demethylation assays

The human AT2-like cell line A549 (CCL-185, ATCC) was grown in Ham’s F12 medium (PAN Biotech, P04-14550) supplemented with 10% fetal bovine serum (FBS, Gibco, 10270106), 1% Glutamax (Gibco, 35050061) and 1% penicillin-streptomycin (Fisher Scientific, 15140122) at 37C in 5% CO_2_ atmosphere, as recommended to preserve the AT2-like phenotype (Cooper et al. 2016). Thus, A549 cells were maintained in Ham’s F12 medium Cells were purchased from the ATCC and routinely tested for Mycoplasma, and throughout the experiments they tested negative. For demethylation assays cells were seeded at 10^3^ cells per cm^2^ in 21 cm^2^ cell-culture treated dishes (Falcon). 48h later cells received 5-Aza-2’-deoxycytidine (0,5µM AZA, SIGMA, A3656) or DMSO and medium was replaced 48h after. 72h after treatment initiation cells were left to recover for 48 hours in complete Ham’s F12 medium without AZA. Each experiment was performed in 3 independent biological replicates. Total RNA and DNA was isolated using AllPrep DNA/RNA Micro Kit (QIAGEN, 80284).

### Validation of LINE demethylation with Mass Array

Demethylation of LINE elements was validated using matrix-assisted time-of-flight mass spectrometry (MassARRAY; Agena Bioscience), a sequencing-independent method. The MassARRAY assay was performed as described previously (Ehrich et al. 2005). Specific primers targeting LINE-1 were used on bisulfite-treated genomic DNA from A549 cells treated with 0,5µM AZA or DMSO (Fw: TTTATATTTTGGTATGATTTTGTAG; Rv: TTTATCACCACCAAACCTACCCT). To quantify the level of methylation of the 3 CpG sites, a DNA methylation standard with defined ratios of in vitro methylated whole genome amplified DNA was included (0, 20, 40, 60, 80 and 100%).

### Gene expression analysis using quantitative PCR

A549 cells were harvested, and RNA and DNA were isolated using the AllPrep DNA/RNA micro kit (QIAGEN) according to the manufacturer’s instructions. 1 microgram of total RNA was reverse transcribed using Revertaid 1st cDNA synthesis kit (Thermo Fisher Scientific) according to the supplier’s protocol. To quantify the expression of IFN pathway genes, real-time PCR was performed with 10 ng of cDNA and gene-specific TaqMan assays as suggested in the manual (RPLP0: Hs00420895_gH; HLA-F: Hs01587837_g1; IL33: Hs04931857_m1; IL6: Hs00174131_m1; IRF5: Hs00158114_m1; IRF9: Hs00196051_m1; MX2: Hs01550814_m1; PSMB8: Hs00544758_m1; PSMB9: Hs00160610_m1; OAS1: Hs00973635_m1; OAS2: Hs00942643_m1; TAP2: Hs00241060_m1). To quantify the expression of other genes, real-time PCR was performed using 10 ng of cDNA, Fast SYBR® Green (ThermoFisher) and specific KiCqStart® SYBR® Green primers (Merck). MicroAmp™ Optical 384-Well Reaction with 10µL reactions were loaded into a QuantStudio™ 5 Real-Time PCR System (Applied Biosystems) and run according to the following recommended program 10 min 55C, 1 min 95C, followed by 40 cycles of 10 s 95C, 1 min 60C. For each biological replicate, all reactions were run in duplicates and the average C_T_ values between duplicates were used for analysis. The fold change in gene expression upon AZA treatment was calculated using the (2^-ΔΔC^_T_) compared to control DMSO treated cells after normalization to RPLP0 expression.

### Epigenetic editing of IRF9 gene

A549 cells were maintained as described above. 19 different sgRNAs targeting promoter (P), or gene body (GB) regions of the human *IRF9* gene were designed using E-CRISP (Heigwer et al. 2014) and CRISPOR (Concordet and Haeussler 2018), and purchased from Integrated DNA Technologies (IDT) as separate oligo strands with overhangs complementary to the BbsI cleavage site. The two strands were annealed at equimolar ratio by heating to 95°C and allowing to cool and ligated into the BbsI-HF-digested empty U6_gRNA vector (a gift from George Church Addgene plasmid #41824, (Mali et al. 2013)), modified to contain two BbsI sites at the gRNA sequence position (Stepper 2020). For targeted DNA demethylation, the plasmid dSPn-VPR-mTET3del1ΔC, consisting of an engineered catalytic domain of mouse TET3 fused to tripartite VP64-p65-Rta (VPR) transcriptional activator was used (Stepper 2020).

For epigenetic editing, A549 cells were seeded into a 6-well plate, 600,000 cells per well to achieve 70% confluency the following day. The cells were transfected using polyethyleneimine (PEI, MW 40,000; 1 mg/ml, Polysciences) using 2 µg of plasmid DNA ad 6 µl of PEI. Several different setups were used. Transfection control contained 5% pEGFP puro (a gift from Michael McVoy, Addgene plasmid #45561(Abbate et al. 2001)), and either (i) 20% pUC19 and 75% empty modified U6_gRNA vector for the mock transfection control; (ii) 20% dSPn-VPR-mTET3del1ΔC and 75% empty modified U6_gRNA vector for the non-targeted control; or (iii) 20% dSPn-VPR-mTET3del1ΔC and 75% pooled IRF9 gRNAs (either mix 1, gRNAs 1-10, targeting the promoter (P) region of *IRF9*, mix 2, gRNAs 11-19, targeting the gene body (GB) region of *IRF9* or a mix containing all 19 gRNAs targeting both P and GB regions of *IRF9*). 24 hours later, the medium was replaced with 2 mL of prewarmed full growth medium containing 2 µg/mL of puromycin and 25 mM of sodium L-ascorbate (Sigma-Aldrich). Puromycin selection was performed for 2 days, by replacing the medium every 24 hours. Upon reaching confluency, cells were transferred to T25 and maintained until Day 10 post transfection in the presence of 25 mM of sodium L-ascorbate. On Day 10 cells were pelleted, flash frozen in liquid nitrogen and stored at −80 C until RNA and DNA isolation.

### Bisulphite amplicon sequencing

Genomic DNA was extracted from 3 independent replicates following the Bio-On-Magnetic-Beads (BOMB) DNA extraction protocol using silica-coated BOMB beads (Oberacker et al. 2019). Extracted DNA was converted using the EZ DNA Methylation Kit (Zymo Research). Bisulphite PCR primers targeting the IRF9 P and GB regions were designed against the human genome assembly 38 (hg38) using the Zymo Research Bisulphite Primer Seeker online design tool (https://zymoresearch.eu/pages/bisulfite-primer-seeker). HotStarTaq plus PCR reagents (Qiagen) were used for PCR amplification. Bisulphite PCR amplicons were purified using the carboxyl-coated BOMB bead Clean Up protocol (Oberacker et al. 2019). Amplicons were phosphorylated with T4 PNK (NEB) and ligated with TruSeq adapters using T4 DNA ligase (NEB). Dual-indexing was carried out with the TruSeq panel of indexing primers in a short-cycle PCR using Luna Universal Probe One-Step RT-qPCR reagents (NEB). The pooled adapter libraries were purified and size-selected using AMPure XP beads (Beckman Coulter) or carboxyl-coated BOMB beads (Oberacker et al. 2019), and quantified with Qubit Fluorometer using dsDNA-HS reagents (Invitrogen). Samples were sequenced by the Cardiff University School of Biosciences Genomics Research Hub on an Illumina MiSeq using Nano flow cells with V2 reagents in paired-end mode with 500 cycles to yield 450 bps forward and 50 bps reverse reads. The sequencing library was loaded at a concentration of 8pM with 25% PhiX DNA added to maintain read complexity.

### Sequence analysis

Analysis of sequencing data was performed as previously described (Stepper et al. 2017). Poor quality reads and adapters were filtered and trimmed with Trim Galore (v.0.6.10; https://www.bioinformatics.babraham.ac.uk/projects/trim_galore/) using default settings. Bismark (v.0.24.0; https://www.bioinformatics.babraham.ac.uk/projects/bismark/) was used to align reads to the human genome (hg38) with default settings and to extract methylation data that were then processed using SeqMonk (v. 1.48.1; https://www.bioinformatics.babraham.ac.uk/projects/seqmonk/) to quantify percentage methylation at each CpG site. This data was further visualized using UCSC Genome Browser custom tracks function. At least 60-fold coverage was achieved for all but one sample.

### Quantification of gene expression after epigenetic editing

The RNA was isolated from 3 independent replicates following the Bio-On-Magnetic-Beads (BOMB) protocol (Oberacker et al. 2019). RevertAid First Strand cDNA Synthesis Kit (Thermo Fisher Scientific) was used for cDNA synthesis following the manufacturer’s instructions. The quantitative PCR reaction mix contained 5 µl of the Luna Universal qPCR Master Mix (2X, New England BioLabs), 1 µl of the forward primer (10 µM), 1 µl of the reversed primer (10 µM), 0.5 µl of 10X SYBR dye and 0.5 µl of nuclease-free water. 10 ng of cDNA was used per reaction and technical triplicates were run for each primer pair. The primers used for qPCR can be seen below. The samples were run in QuantStudio™ 5 Real-Time PCR System using the following program: 50 °C for 2 mins, 95°C for 10 mins, 40 cycles at 95 °C for 15 secs and 60°C for 1 min, followed by 95 C for 15 secs, 60°C for 1 min and 95°C for 0.1 sec. For each run, the melting curves were collected at the end of the experiment to assess the amplification specificity. The qPCR data were normalized to the housekeeping gene REEP5. To calculate the expression change of a target gene, 2^-ΔΔC^_T_ values were normalized to the mock transfection control (pUC19). Statistical analysis was performed with GraphPad Prism software, version 8.0.1 or with R. The significance level was set at 0.05 unless otherwise indicated. The number of replicates and the statistical test used are described in the figure legends for each of the panels.

An unpaired nonparametric t-test (Mann-Whitney test, GraphPad Prism software, version 8.0.1) was used to compare lung function (FEV_1_ and FEV_1_/FVC values) between control and COPD donors. Non-parametric one-way ANOVA (Kruskal-Wallis test, GraphPad Prism software, version 8.0.1) followed by correction for multiple comparisons using Dunn’s test was used to analyze the patient metadata of the three groups studied (control, COPD I, and COPD II-IV).

RT-qPCR analysis of the 5-aza demethylation assay was performed by paired t-test, FDR-corrected using two-stage linear step-up procedure of the Benjamini, Krieger, and Yekutieli, with Q = 5%. Each row was analyzed individually, without assuming a consistent SD. *p<0.05; **p<0.005; *** p < 0.0005

For epigenetic editing, statistical significance of data was analyzed using the Kruskal-Wallis multiple comparison test, followed by Dunn’s post-hoc analysis comparing each sample against the NTC (total 3 comparisons), and adjusted p-values using the Benjamini-Hochberg correction method (*p < 0.05; **p < 0.01; ***p < 0.001, ****p < 0.0001).

## Data availability

The WGBS and RNA-seq data generated in this study have been deposited at the European Genome-phenome Archive (EGA, https://ega-archive.org), which is hosted by the EBI and the CRG. The access to the patient data is controlled by the data access committee.

- RNA-seq data: EGAS00001007387
- T-WGBS DNA methylation data: GAS00001007386

## Author contributions

RZJ, US & MLP contributed to the design and conception of the study. MLP, SP, DP, AB and DA performed experiments with the help from RL, RT, MR and EE. US performed bioinformatic analysis and integration of the RNA-seq and WGBS data. TMu, MS, FH, HW provided lung tissue and patient data. AW performed the pathological analysis of the H&E lung specimens. CPH determined the emphysema score index of the patients. HS, JH, DWy, TPJ, CI, BB, and CP provided critical input, analysis software and/or materials. RZJ, US and MLP wrote the manuscript with the input from all authors. All authors contributed to scientific discussions and approved the final version of the manuscript.

## Acknowledgments

We would like to thank Lung Biobank (Heidelberg, Germany) – a member of the Biomaterial bank Heidelberg (BMBH), the tissue bank of the National Center for Tumor Diseases (NCT) and the Biobank platform of the German Center for Lung Research (DZL) for providing Biomaterials and Data. We also thank Christa Stolp for help with collecting primary material. We acknowledge excellent sequencing service and helpful discussions from the Genomics core facility (GeneCore, EMBL, Germany) for RNA-seq, from NGX Bio (San Francisco, USA) for T-WGBS sequencing, and from Cardiff University School of Biosciences Genomics Research Hub for bisulfite amplicon sequencing, as well as support from the ZMBH imaging facility (Heidelberg, Germany) for immunofluorescence. We thank Morphisto GmbH (Frankfurt, Germany) for excellent histological service, Peter Stepper (Stuttgart University, Germany) for providing dSPn-VPR-mTET3del1ΔC vector and Olivier Mücke (DKFZ, Germany) for running the Mass array analysis. We also thank Christian Tidona (BioMed X Innovation Center) and Markus Koester (Boehringer Ingelheim) for helpful project discussions.

---

**Figure S1.**
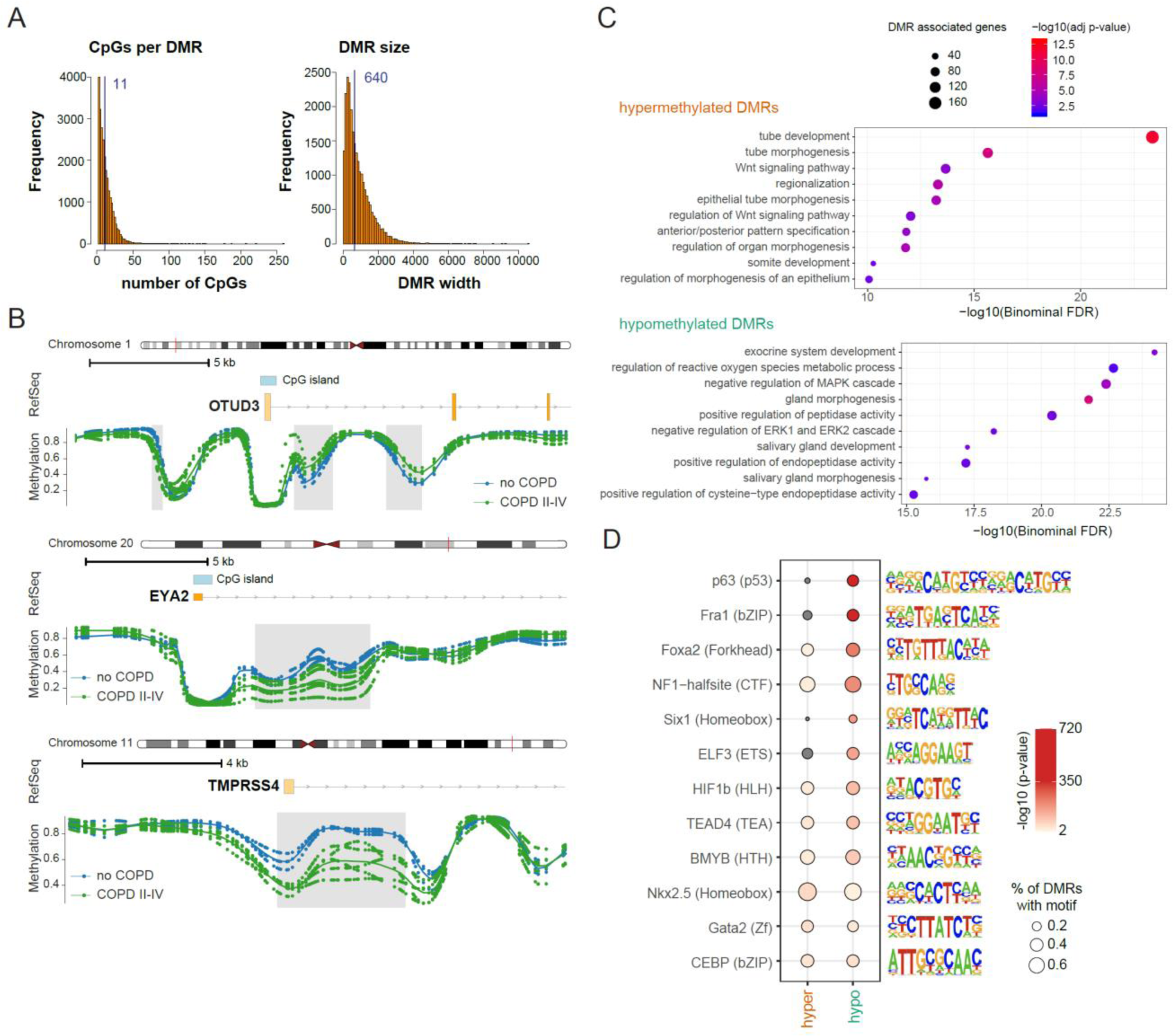
Genome-wide DNA methylation changes occur at regulatory regions in AT2 cells during COPD (supporting information for Figure 2). **A**. Number of CpG sites (left panel) and the width of DMRs (right panel) identified between no COPD and COPD II-IV. Median values are indicated in the histograms in dark blue. **B**. Functional annotation of genes located next to hypermethylated (top) and hypomethylated (bottom) DMRs using GREAT. Hits were sorted according to the binominal adjusted p-value and the top 10 hits are shown. The adjusted p-value is indicated by the color code and the number of DMR associated genes is indicated by the node size. **C**. Detailed view of DMRs showing the methylation profiles of no COPD (n=3) and COPD II-IV (n=5) samples at the indicated genomic regions. DMR locations are highlighted as grey boxes. **D**. Transcription factor motif enrichment in hypermethylated (left) and hypomethylated (right) DMRs. The top motif of each transcription factor family (in brackets) is shown. The node size indicates the percentage of DMRs containing the respective motif and the color represents the p-value of the enrichment analysis.

**Figure S2.**
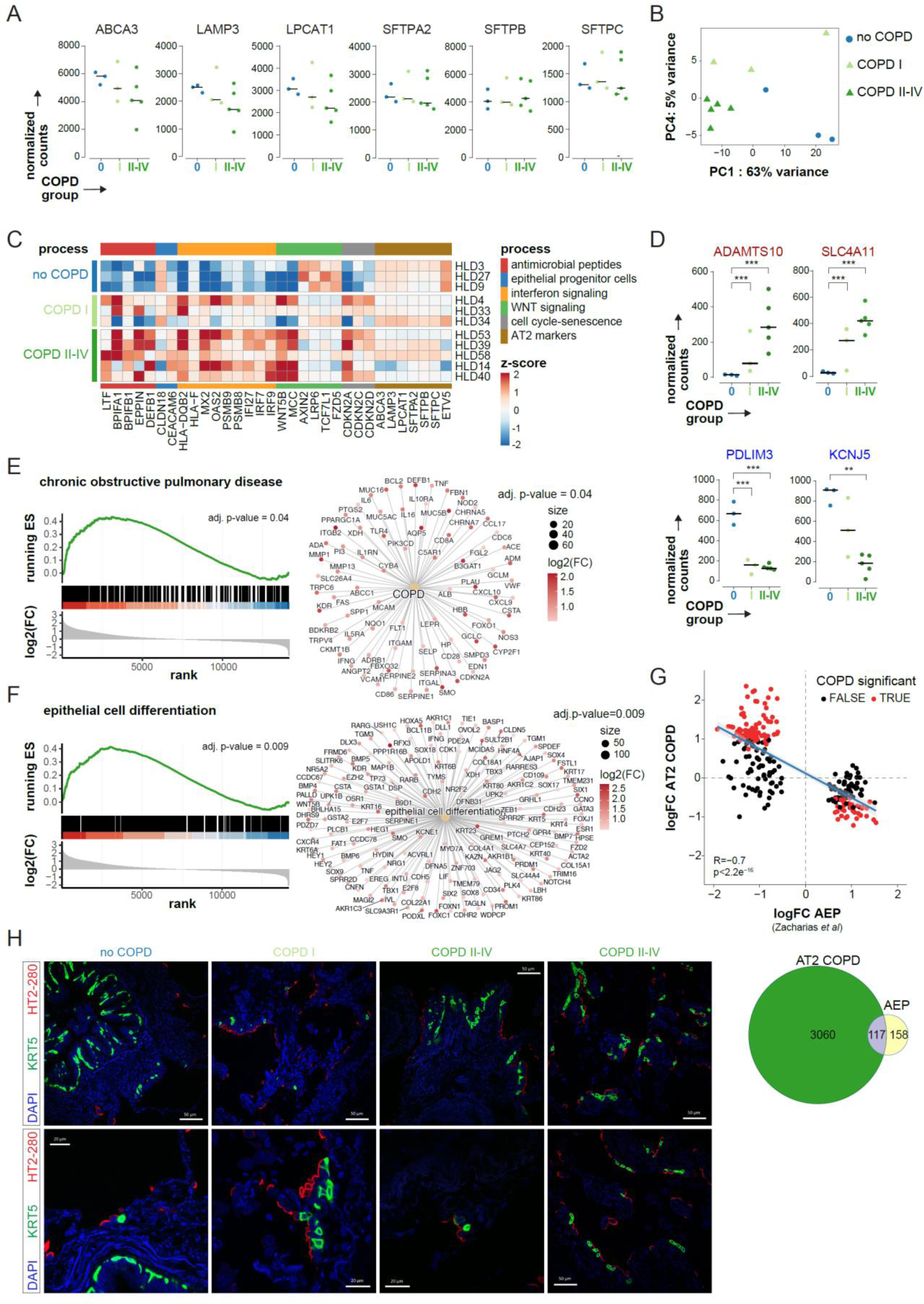
AT2 transcriptome is altered in COPD as disease progresses (Supporting information for Figure 3) **A**. Normalized read counts from RNA-seq data for AT2 specific genes in sorted AT2 cells from each donor (dots). Data points represent normalized counts from no COPD (blue), COPD I (light green) and COPD II-IV (dark green). The group median is shown as a black bar. **B**. Principal component analysis (PCA) of the 500 most variable genes. Percentage indicates proportion of variance explained by PC1 and PC4. COPD I and COPD II-IV samples are represented in light and dark green triangles, respectively, and no COPD samples as blue circles. **C**. Heatmap showing expression changes of selected genes across all samples. Genes associated with selected processes are shown. The expression deviation from the average expression across all samples (log(norm. expression/average expression) is indicated by the color code. **D**. Top upregulated (top) and top downregulated (bottom) genes in AT2 cells from COPD patients. Normalized read counts from RNA-seq data for the specified genes in each donor (dots). Data points represent normalized counts from no COPD (blue), COPD I (light green) and COPD II-IV (dark green). The group median is shown as a black bar. FDRs were calculated using DESeq2, which uses a negative binominal GLM (generalized linear model) and Wald statistics. **: FDR <0.01, ***: FDR <0.001. **E-F**. Gene set enrichment analysis (GSEA) results of the gene expression of AT2 cells from COPD II-IV vs no COPD donors. ES, enrichment score; NES, normalized enrichment score; FDR, false discovery rate. GSEA of genes associated with (**E**) chronic obstructive pulmonary diseases (Diseases Ontology ID: 3083) or (**F**) epithelial cell differentiation (Gene Ontology: GO:0030855). Genes were sorted based on the log2(fold-change) in COPD II-IV (bottom panel). The p-value of the GSEA is indicated in the top right corner. Gene network plot of the respective terms show genes associated with the term and driving the enrichment score (leading edge). The beige nodes symbolize enriched terms, with the size of the node indicating the number of associated genes. The lines connecting the nodes denote the specific genes associated with each term. The color of the gene nodes signifies the expression change in severe COPD compared to no COPD. **G.** Top, scatter plot showing genes differentially expressed in alveolar epithelial progenitor (AEP) cells from (Zacharias et al. 2018) compared to AT2 (Zacharias AEP) and their correlation to AT2 cells in COPD II-IV compared to AT2 from no COPD donors from our study (AT2 COPD). DEG in AEP were determined with the same pipeline and cutoffs (red dots; FDR of 10% and | log2(fold-change) | > 0.5) as used for RNA-seq of no COPD vs COPD II-IV (see Methods section). DEGs in COPD II-IV are highlighted as red dots. The blue diagonal represents the linear regression between log2(fold-changes) in COPD II-IV against no COPD and AEP against AT2. Shaded areas are confidence intervals of the correlation coefficient at 95%. *P*-value was derived from linear regression analysis and the Pearson correlation coefficient (*R*) is indicated. Bottom, Venn diagram indicating the overlap of DEG in AT2 COPD and AEP cells from (Zacharias et al. 2018) compared to AT2 cells. **H.** Representative immunofluorescence staining images of HT2-280 and KRT5 expression in FFPE human lung tissue slices from no COPD, COPD I and COPD II-IV donors. Slides were counterstained with DAPI, scale bars = 50µm or 20µm as displayed in images.

**Figure S3.**
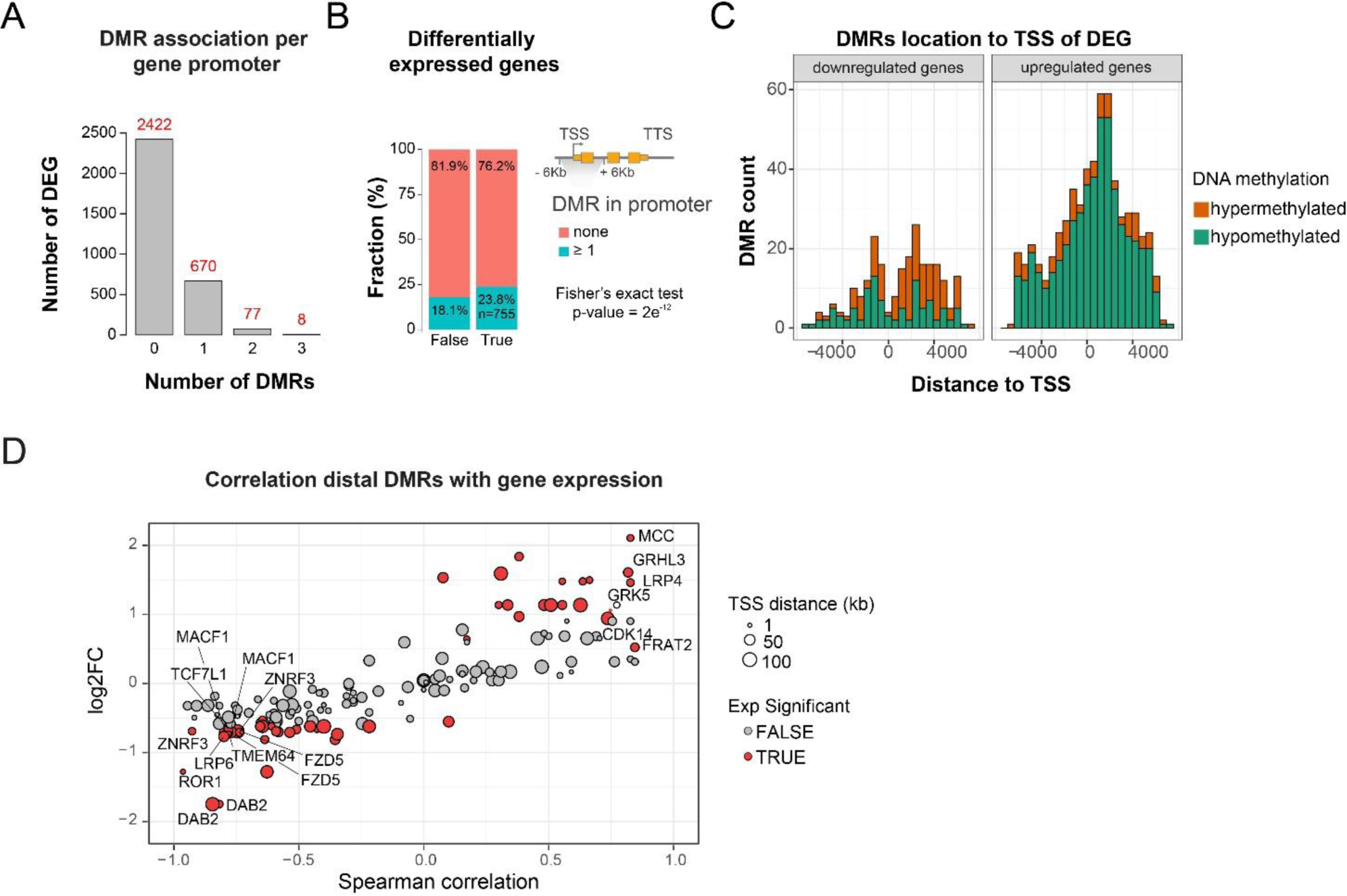
Integrated analysis reveals epigenetic regulation of key pathways in COPD. (Supporting information for Figure 4). **A.** DMRs within ± 6 kb from TSS of DEG were assigned to their corresponding gene. **B**. Fraction of genes associated with at least one DMR in the promoter proximity (+/− 6 kb from TSS, blue) of non-DEG (left, 18.1%) or DEG (right, 23.8%). DMRs are significantly enriched at DEGs (Fisher’s exact test p-value=2e^-12^). **C**. Location of hyper- and hypomethylated DMRs relative to the TSS of DEGs in downregulated (left) and upregulated (right) genes. **D**. Scatter plot showing distal DMR-DEG pairs associated with Wnt-signaling. Pairs were extracted from GREAT analysis (hypermethylated, DMR-DEG distance < 100 kb; **Fig. S1C**). Y-axis represents the expression log2 fold-change in severe COPD of the respective gene and the x-axis denotes the Spearman correlation of the DEG-DMR pair. The node size indicates the distance of the DMR to the TSS. DEGs are highlighted by red color.

**Figure S4.**
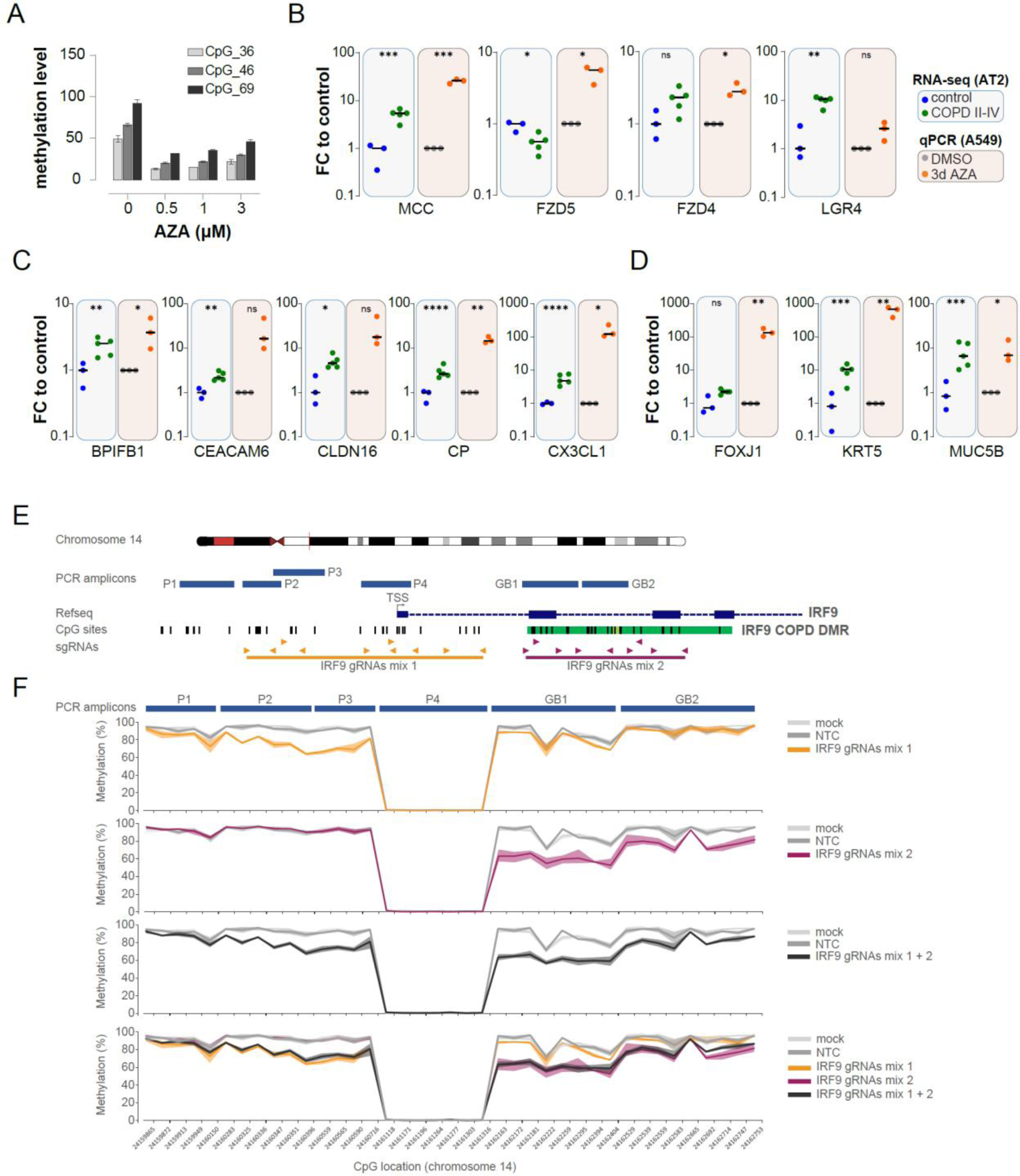
Epigenetic regulation of gene expression in AT2 and A549 cells. **A**. LINE methylation levels in A549 cells treated with the indicated amounts of 5-Aza-2’- deoxycytidine (AZA) and measured at three CpG sites by Mass Array. **B-D**. Fold-change in gene expression of selected genes in AT2 cells in COPD (RNA-seq) and in A549 cells treated with 0.5µM AZA (RT-qPCR) compared to control samples. Genes were selected based on the presence of a DMR and deregulation in AT2 cells in COPD. Left, RNA-seq data from AT2 cells (no COPD, blue, n=3; COPD II-IV, green, n=5), right, A549 treated with AZA (orange, n=3) compared to control DMSO treated cells (grey, n=3). The group median is shown as a black bar. Selected genes from the WNT/β-catenin pathway (**B**), antimicrobial peptides, alveolar progenitor (**C**) and airway epithelial markers (**D**) are shown. RNA-seq FDRs were calculated using DESeq2 with negative binomial generalized linear model (GLM) and Wald statistics. *: FDR <0.05; **: FDR <0.01. Gene expression was measured by RT-qPCR using DMSO treatment as control (grey) and RPLP0 as housekeeping gene. For A549 samples, each point represents the mean of two technical replicates, and bars represent the median of 3 independent experiments (n=3). Statistical analysis was performed by paired t-test, FDR-corrected using the Benjamini, Krieger, and Yekutieli two-stage set-up method. *p-value < 0.05; **p-value < 0.01. **E**. Graphical representation of the IRF9 locus. In dark blue is the IRF9 coding region, with introns and exons represented by dashed and solid lines, respectively. At the bottom, arrows represent individual gRNAs targeting the IRF9 promoter (orange) and gene body (magenta) regions, with overlapping lines representing the groups of gRNAs that comprise mix 1 and mix 2, respectively. Bisulfite-PCR amplicons targeting IRF9 (blue bars) are depicted as blue boxes, with P1-4 targeting the IRF9 promoter region, and GB1-2 targeting the gene body region. Individual CpG sites are depicted by black bars below the IRF9 coding region, with the light and dark green overlapping region displaying the core and extended differentially methylated regions, respectively, identified in the AT2 COPD TWGBS data. Genomic positions were extracted from human genome assembly 38 (hg38) using the UCSC genome. **F**. CpG-methylation percentage at individual IRF9 CpG sites in epi-edited A549 cells. The percentage CpG-methylation for mix 1, 2, and 1+2 (orange, magenta, and dark grey) transfected samples are plotted separately against the pUC19 mock (mock) and non-targeted (NTC) transfection controls. For each sample, the opaque line plots the mean value, and the faint surrounding area plots the observed range across repeats. The bisulfite PCR targets are displayed across the top, showing which CpG-sites are sequenced by each target, and how these sites relate to the genomic locations of the sequencing targets seen in panel.

